# Recombination and sterility in inversion homo- and heterokaryotypes under a general counting model of chiasma interference

**DOI:** 10.1101/2020.03.28.013144

**Authors:** Øystein Kapperud

## Abstract

It has long been known that chiasmata are not independently generated along the chromosome, a phenomenon known as *chiasma interference*. In this paper, I suggest a model of chiasma interference that generalizes the *Poisson model*, the *counting model*, the *Poisson-skip model* and the *two-pathway counting model* into a single framework, and use it to derive infinite series expressions for the sterility and recombination pattern probabilities in inversion homo- and heterokaryotypes, and a closed-form expression for the special case of the two-pathway counting model in homokaryotypes.

## 1 Introduction

Meiotic recombination reshuffles the genetic material before reproduction, and may thus serve to avoid the problem of having slightly deleterious mutations accumulate through their association with favored ones (Muller 1964). Although models of multilocus evolution typically assume that the foci of recombination are independently distributed along the chromosome, in reality it has long been known that such *chiasmata* are subject to spatial *interference* (Sturtevant 1913, 1915, Muller 1916, Berchowitz and Copenhaver 2010). In 1993, Foss et al. suggested the *counting model* of chiasma interference, in which the genetic distances between neighbouring chiasmata are generated by summing a given number of exponential distributions. This model later spawned the *Poisson-skip model* of Lange et al. (1997), in which the number of intervening exponential distributions is drawn from a given probability distribution, and the *two-pathway counting model* of Copenhaver et al. (2002), which is identical to Foss et al.’s model except that it allows for a second pathway of non-interfering chiasmata. These models, which I will refer to collectively as the *counting models*, have been used to generate expressions of recombination pattern probabilities which fit well with data for inversion homokaryotypes from a wide range of organisms (Foss et al. 1993, Zhao et al. 1995a, Lande and Stahl 1993, Lange et al. 1997, Copenhaver et al. 2002).

Chromosomal inversions partly suppress recombination in heterokaryotypes (e.g. Coyne et al. 1991, 1993, Navarro and Ruiz 1997, Jaarola et al. 1998) and are often found to be linked to loci involved in differentiation of alternative mating strategies (Tuttle et al. 2016, Lamichhaney et al. 2016, Wang et al. 2013), local adaptation (Etges and Levitan 2004, Sinclair-Waters et al. 2018) and pre- and postzygotic reproductive isolation between species or subpopulations (Noor et al. 2001, Feder et al. 2003, Ayala et al. 2013, Poelstra et al. 2014). Although the effects of chiasma interference on underdominance and recombination in inversion heterokaryotypes have been recognized (Navarro et al. 1997), general mathematical expressions for an indefinite number of loci are currently lacking. Accordingly, although the evolution of chromosomal inversions is a common target for modelers (e.g. Charlesworth and Charlesworth 1973, Tricket and Butlin 1994, Navarro and Barton 2003, Feder and Nosil 2009, Dagilis and Kirkpatrick 2016), the potential effects of chiasma interference is to my knowledge invariably ignored.

In this paper, I will present a *general counting model* of chiasma interference, which expands the *Poisson-skip model* to allow for a second pathway of non-interfering chiasmata. Since this model can be reduced to each of the other counting models in special cases, it provides a single unifying framework for investigating chiasma interference. I will here use it to derive exact infinite series expressions for gamete proportions and sterility in inversion homo- and heterokaryotypes for an indefinite number of loci at any location on the chromosome, and a closed form expression for the gamete proportions in homokaryotypes under the *two-pathway counting model*. I single out for special treatment the case of paracentric inversions in Drosophila females, for which meiotic peculiarities complicates the mathematics (Sturtevant and Beadle 1936, Navarro et al. 1997)

### 1.1 General terminology

#### 1.1.1 Recombination

Since discussions of recombination and related issues risk being obfuscated by inconsistent and ambiguous terminology, I will begin by carefully laying out my own. A *tetrad* is a bundle of four *chromatids*, originating as a duplication of each of the two *parental homologues*, with one *pair* of chromatids denoted *sister chromatids* if they are derived from the same homologue and *non-sister chromatids* if they are not. Exchanges of genetic material – or *crossing over –* occur as a series of *chiasmata* (singular *chiasma*) or *chiasma events* distributed along the tetrad according to a model of *chiasma interference*, so that each such event *involves* two *non-sister chromatids*, or, equivalently, one out of the four possible *non-sister chromatid pairs*, chosen according to a model of *chromatid interference*.^1^ In accordance with the lack of consistent evidence to the contrary (Zhao et al. 1995b), I will here assume that each non-sister chromatid pair has an equal chance of being involved in each chiasma event, i.e. I assume no *chromatid* interference (not to be confused with *chiasma* interference). The number of chiasma events in an interval 𝕚 will be denoted *X*_𝕚_, so that the *genetic length* of 𝕚 in Morgans is given by 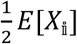. In the following, the terms *distance* and *length* always refer to genetic (as opposed to physical) distance, unless otherwise noted.

A chromatid is said to *be recombinant* or *show recombination* in a given interval if it is *involved* in an odd number of chiasma events within that interval (note that this is not the same as saying that an odd number of chiasma events occur within the interval), and to *be non-recombinant* or show *non-recombination* in the opposite case. A *recombination pattern* is a set of Boolean variables that represent the *recombination status* – recombination (1) or non-recombination (0) – in each of the 𝔫 adjoining and non-overlapping intervals, so that if the set of all intervals – the *region of interest* – is {𝕚_0_, 𝕚_1_, 𝕚_2_ … 𝕚_𝔫−1_}, then we can denote a recombination pattern *r* as

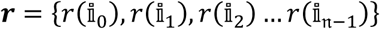

where

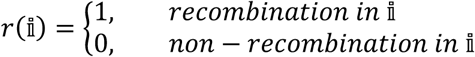

Accordingly, a chromatid is said to *show* or *have* recombination pattern ***r*** if the presence or absence of recombination in all the intervals of the region of interest correspond to ***r***. I will use the random vector ***R*** = {*R*(𝕚_0_), *R*(𝕚_1_), *R*(𝕚_2_) … *R*(𝕚_𝔫−1_)} to represent the recombination pattern of a randomly chosen chromatid, so that I can write e.g. “the probability of observing a chromatid with recombination pattern ***r*** is *p*” as

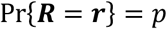

or “the probability of observing recombination status *r*(𝕚) in interval 𝕚 is *p*” as

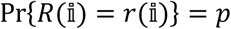

or “the probability of observing recombination in interval 𝕚 is *p*” as

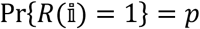

For notational simplicity, I will sometimes use the shorthand forms *R*_*k*_ = *R*(𝕚_*k*_) and *r*_*k*_ = *r*(𝕚_*k*_) when discussing recombination for a set of indexed intervals.

I will assign *directionality* to the region of interest by denoting interval 𝕚_0_ as the *the leftmost interval* and 𝕚_𝔫−1_ as the *rightmost interval*. More generally, I will say that any interval 𝕚_*x*_ is positioned to the *left* of interval 𝕚_*y*_ if *x* < *y*, and to *the right* of 𝕚_*y*_ if *x* > *y*. The directionality also applied to the boundaries between the intervals and the chiasmata within the intervals, so that I for instance can say that a chiasma in interval 𝕚_*x*_ occurs to the right of another chiasma in the same interval if it is positioned closer to the right boundary of 𝕚_*x*_ (which is equivalent to the left boundary of 𝕚_*x*+1_).

#### 1.1.2 Chromosomal inversions

It is common to distinguish between *pericentric inversions*, which include the centromere, and *paracentric inversions*, which do not. In individuals heterozygous for both of these types of inversion, crossing over within the inverted region results in a proportion of *unbalanced gametes* – i.e. gametes that do not have the full set of genes. In the following, all such gametes are assumed to produce inviable zygotes, and the proportion of such gametes for any given individual or genotype will be referred to as that individual or genotype’s *sterility*. For pericentric inversion heterokaryotypes, a chromatid is, as illustrated in figure 4, unbalanced if and only if it shows recombination in the inverted region, so that the sterility is given by the recombination rate of the inverted region. For paracentric inversions, different number of chiasmata in the inverted and proximal regions produce different proportions of tetrads with five different possible *configurations* (figures 1, 4, 6, 7 and 8). In the following, a *no bridge* tetrad is a tetrad that do not form chromosome bridges at either anaphase I or II, a *single anaphase I bridge* tetrad is a tetrad in which one chromatid pair form a chromosome bridge at anaphase I, a *double anaphase I bridge* tetrad is a tetrad in which two chromatid pairs (i.e. all four chromatids) form anaphase I bridges, and a *single* and *double anaphase II bridge* tetrad is the same for anaphase II. I include in theorem 6 a to my knowledge unprecedented expression for the proportions of these five configurations given the number of chiasma events in the inverted and proximal regions. The chromatids involved in a chromosome bridge break randomly at the respective anaphase, and so become unbalanced in the sense defined above.

**Figure 1:**
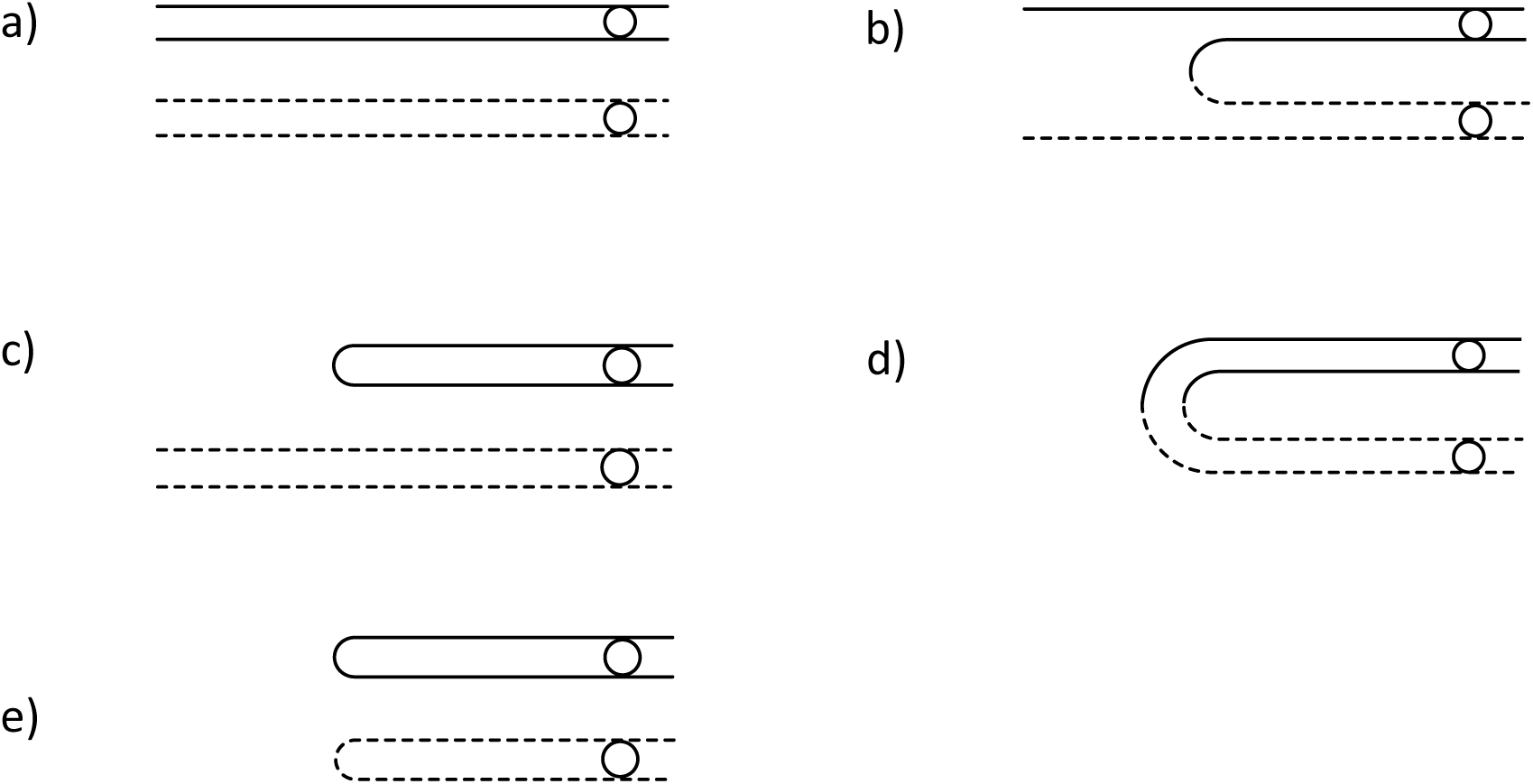
The five types of tetrad configurations. a) no bridge, b) single anaphase I bridge, c) single anaphase II bridge, d) double anaphase I bridge, e) double anaphase II bridge. Acentric fragments (see figure 6) are not shown. Chromatids involved in a bridge break randomly at the given anaphase, and are necessarily unbalanced.

**Figure 2:**
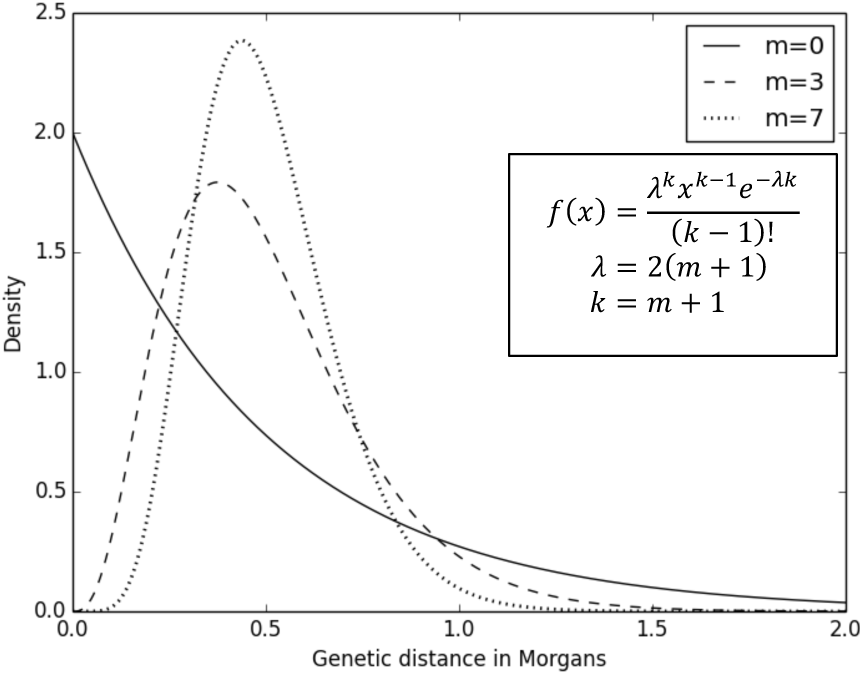
The distribution of distances between chiasma events under the pure counting model with m=0,3,7. Note that all three curves by definition have a mean of 0.5 Morgans, and that the probability of observing two closely spaced chiasma events is smaller for higher m. Since the distances are given by the sum of m+1 (exponential) random variables with rate 2(m+1), it follows from the central limit theorem and the law of large numbers that the distribution approaches an increasingly narrow normal distribution for higher m. In the limit when m approaches infinity, all chiasma events are spaced exactly 0.5 Morgans apart.

**Figure 3:**
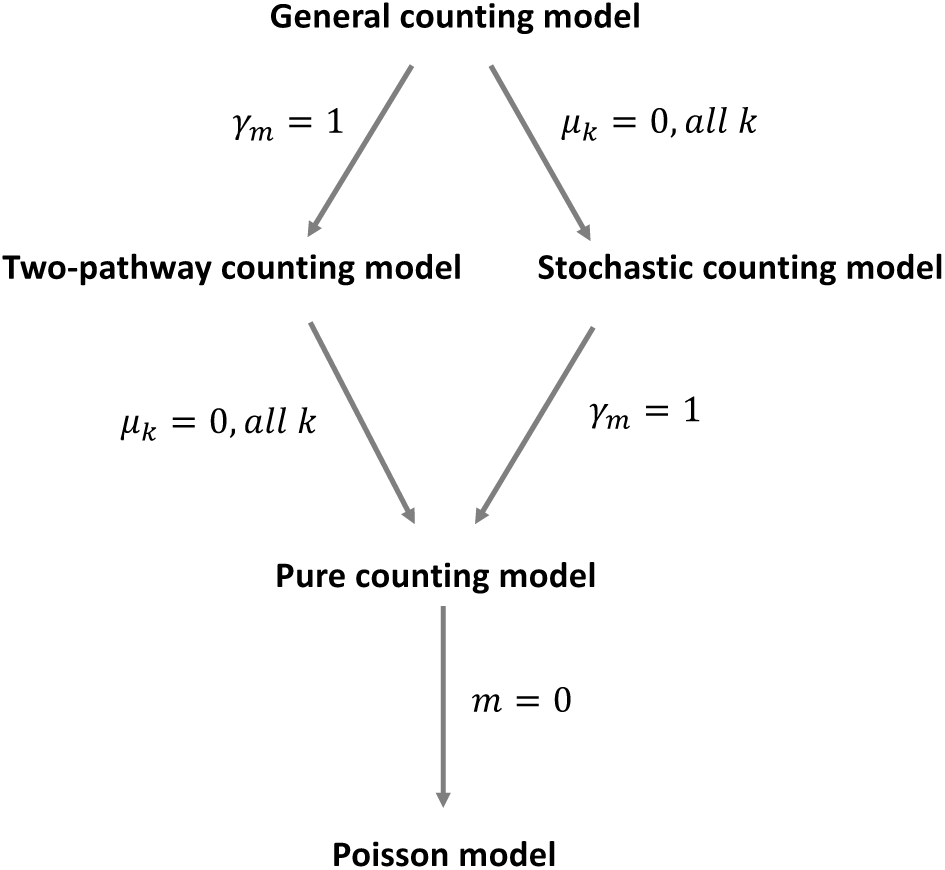
The hierarchy of counting models.

**Figure 4:**
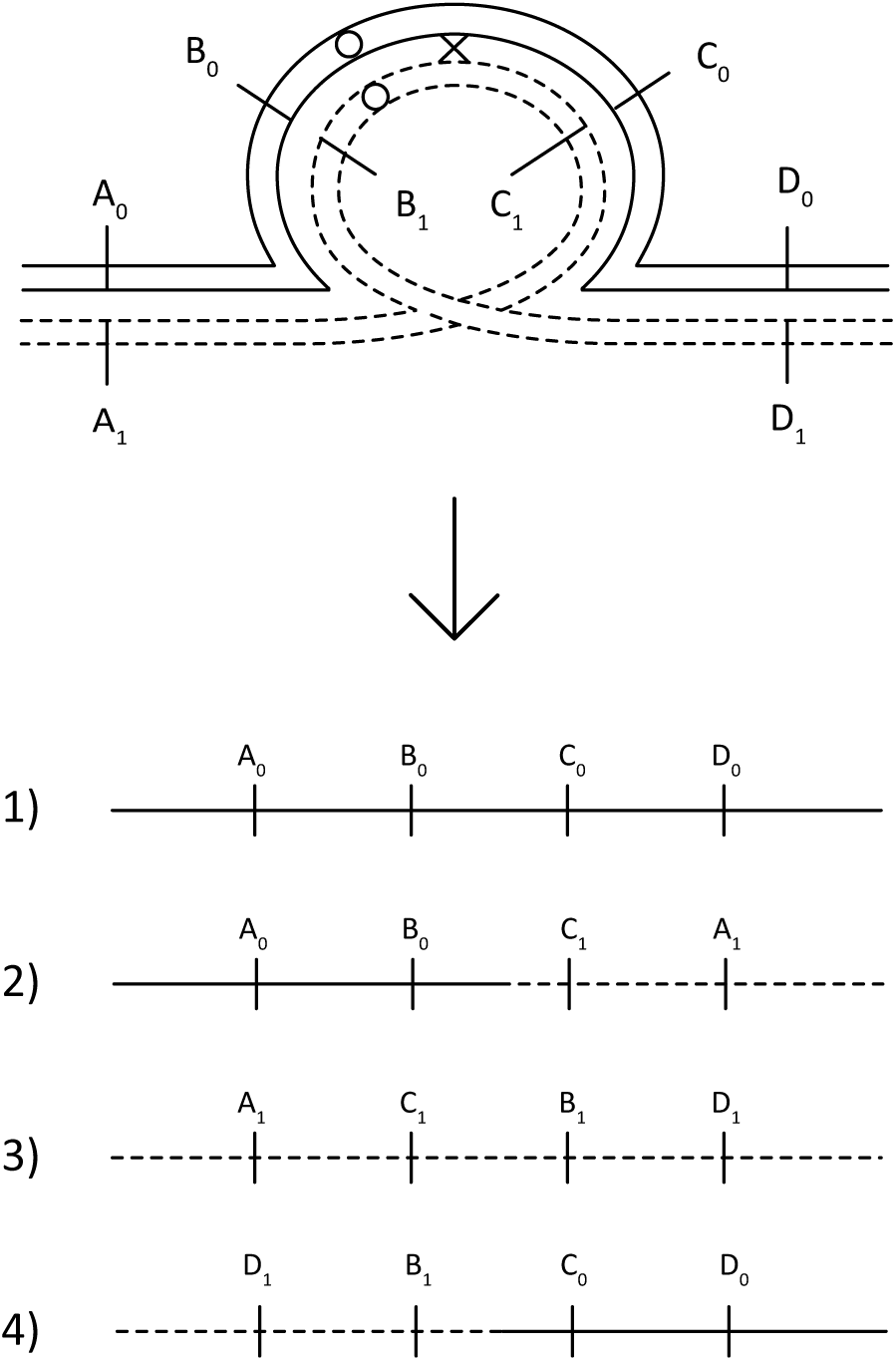
An inversion loop for a pericentric inversion. The lines with the same line type (dotted or dashed) are sister chromatids, and the circles represent the centromere. The X symbolizes a chiasmata, which involves the chromatids touched by the upper and lower arms of the X. Gametes that show recombination in the interval comprising the full inverted region are unbalanced, as illustrated here by gamete 2) and 4), which lack locus D and A, respectively.

**Figure 5:**
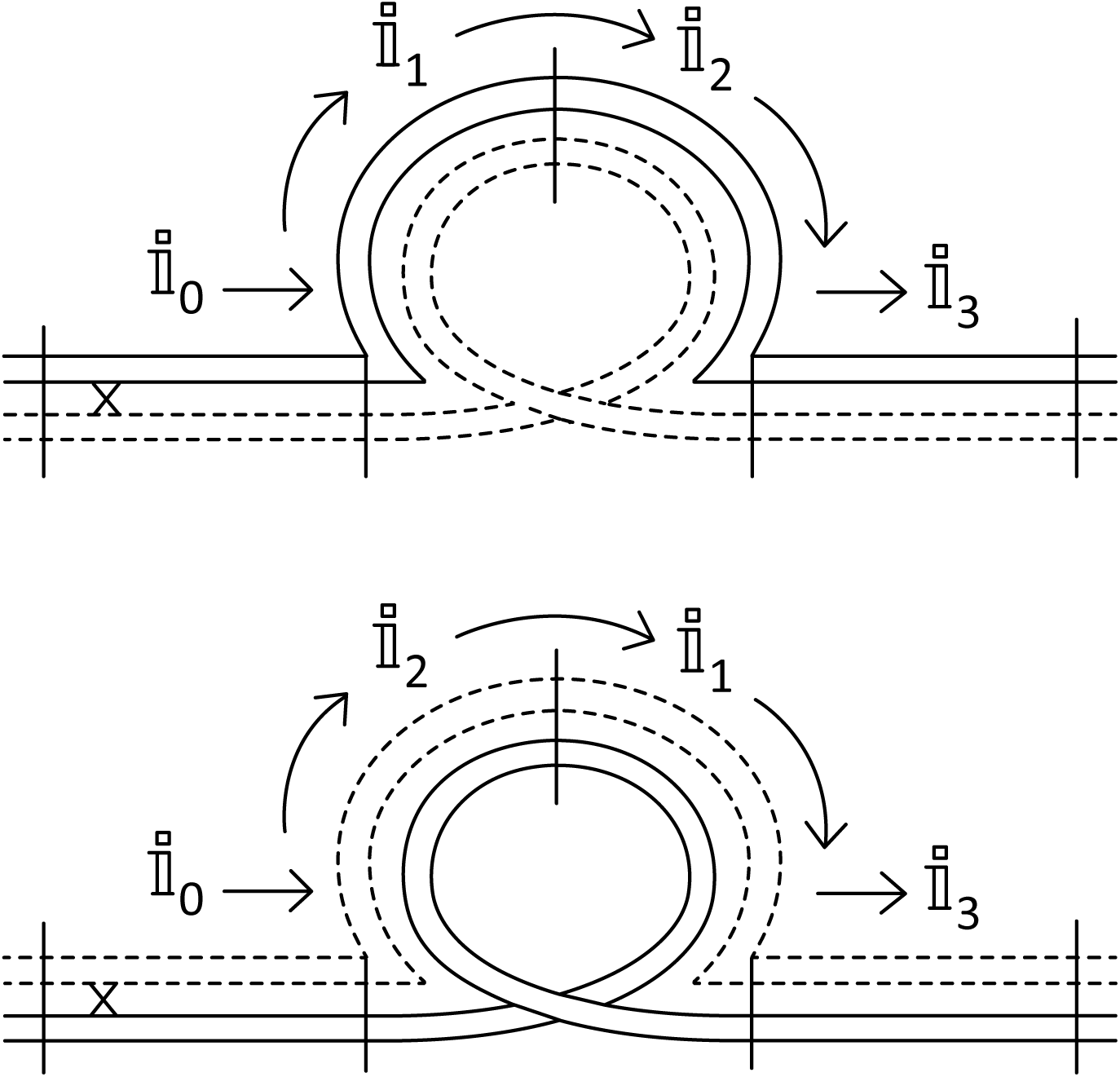
The problem of interference across breakpoint boundaries. Does the interference signal from a chiasma event (the x) in 𝕚_0_ go through 𝕚_1_ (top) or 𝕚_2_ (bottom)? Or is it blocked by the breakpoint boundaries?

**Figure 6:**
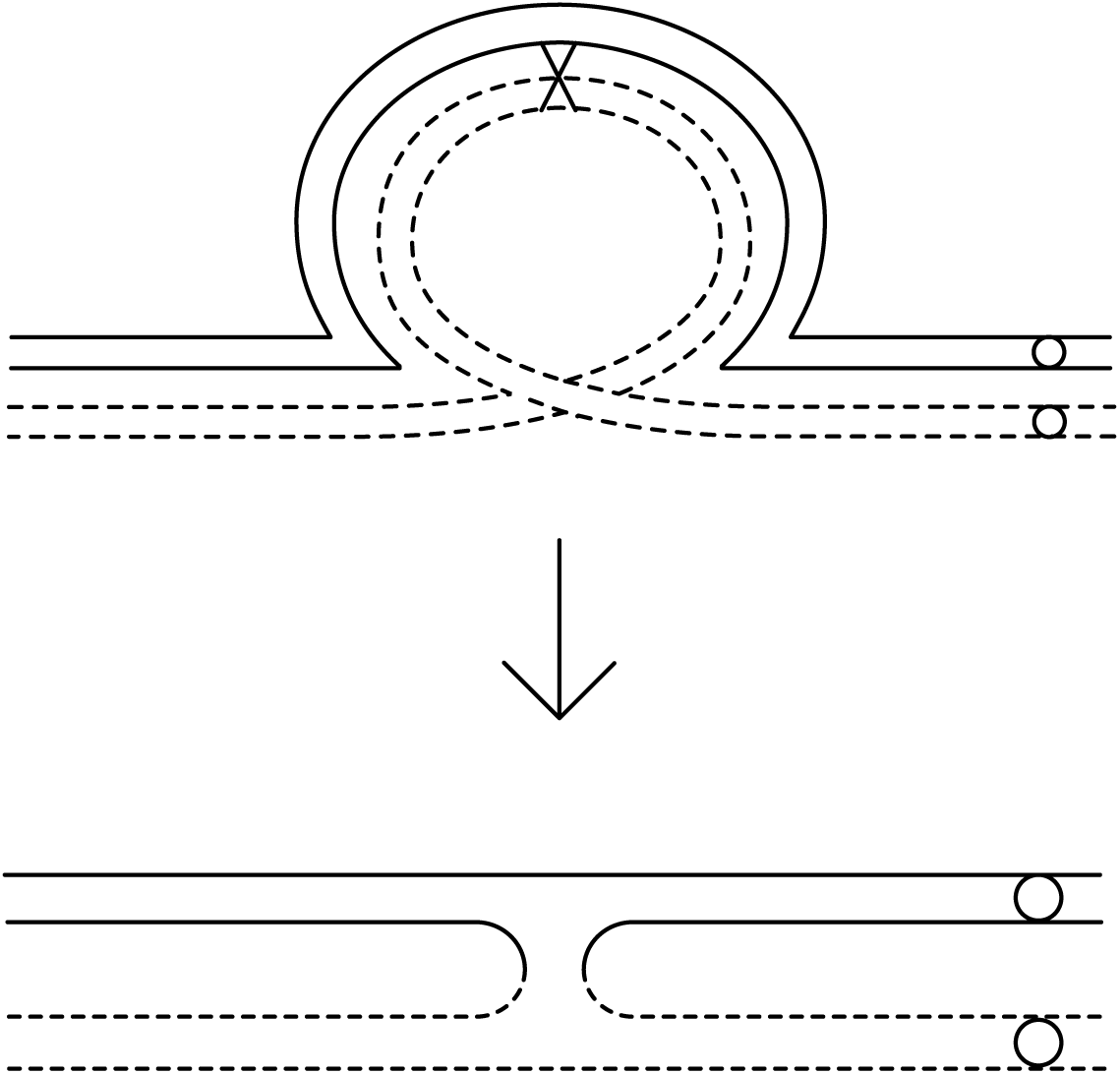
An inversion loop for a paracentric inversion. A single chiasma in the inverted region creates a tetrad with an anaphase I bridge and an acentric fragment (meaning that it is not connected to the centromere) which fails to segregate.

**Figure 7:**
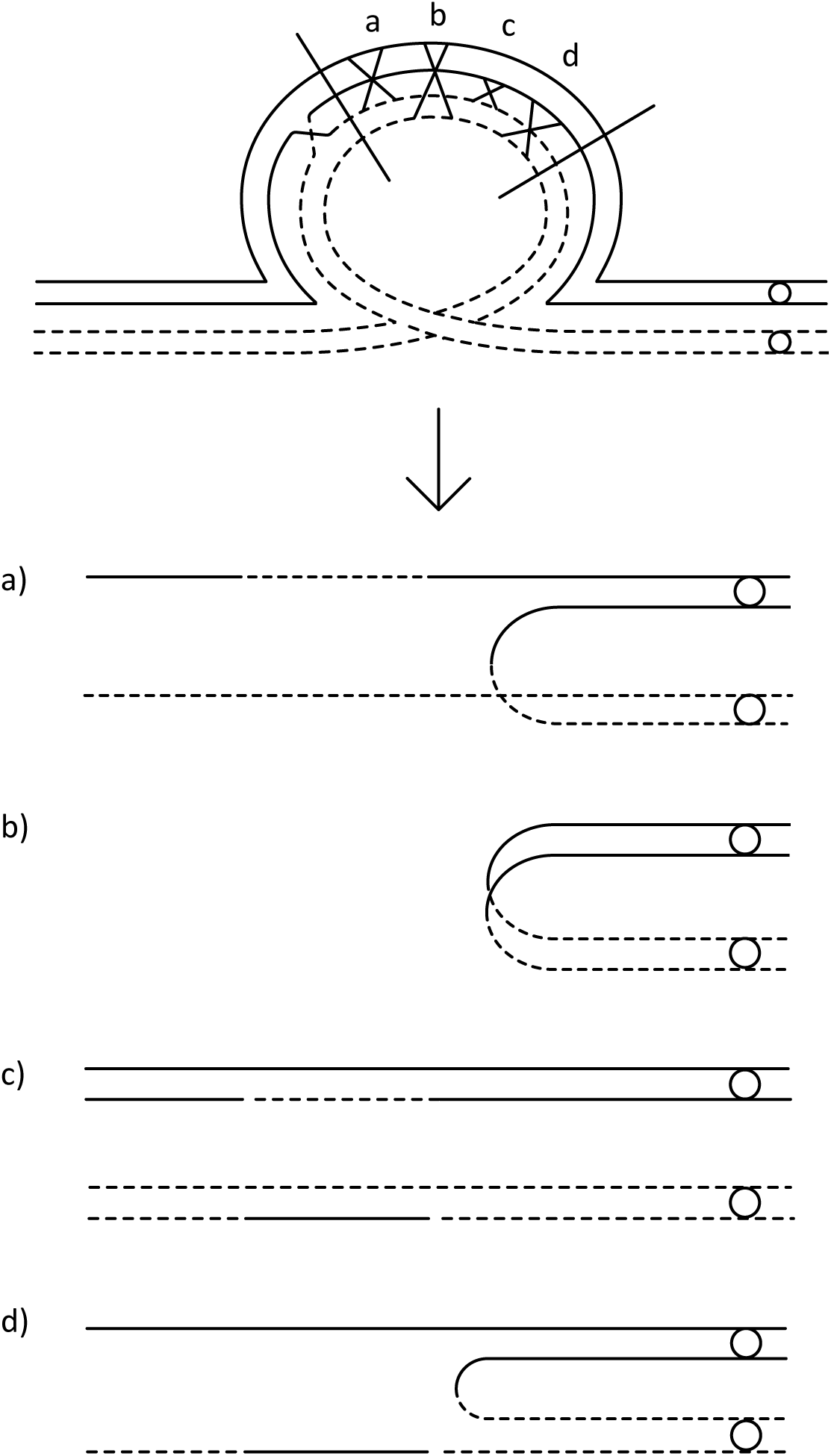
The leftmost X represent an anaphase I bridge, and the X’s marked a, b, c, d represent the four possible non-sister chromatid pair involvements in a single chiasma event. The resulting configurations for each pair is shown below. The figure shows that Pr{Z(x_1_ + 1,0) = 0|Z(x_1_, 0) = 1} = 1/4, Pr{Z(x_1_ + 1,0) = 1|Z(x_1_, 0) = 1} = 1/2, and Pr{Z(x_1_ + 1,0) = 3|Z(x_1_, 0) = 1} = 1/4, which gives the second row (zero-indexed row 1) in matrix **T**_𝕙_. The remaining rows are found in the same manner.

**Figure 8:**
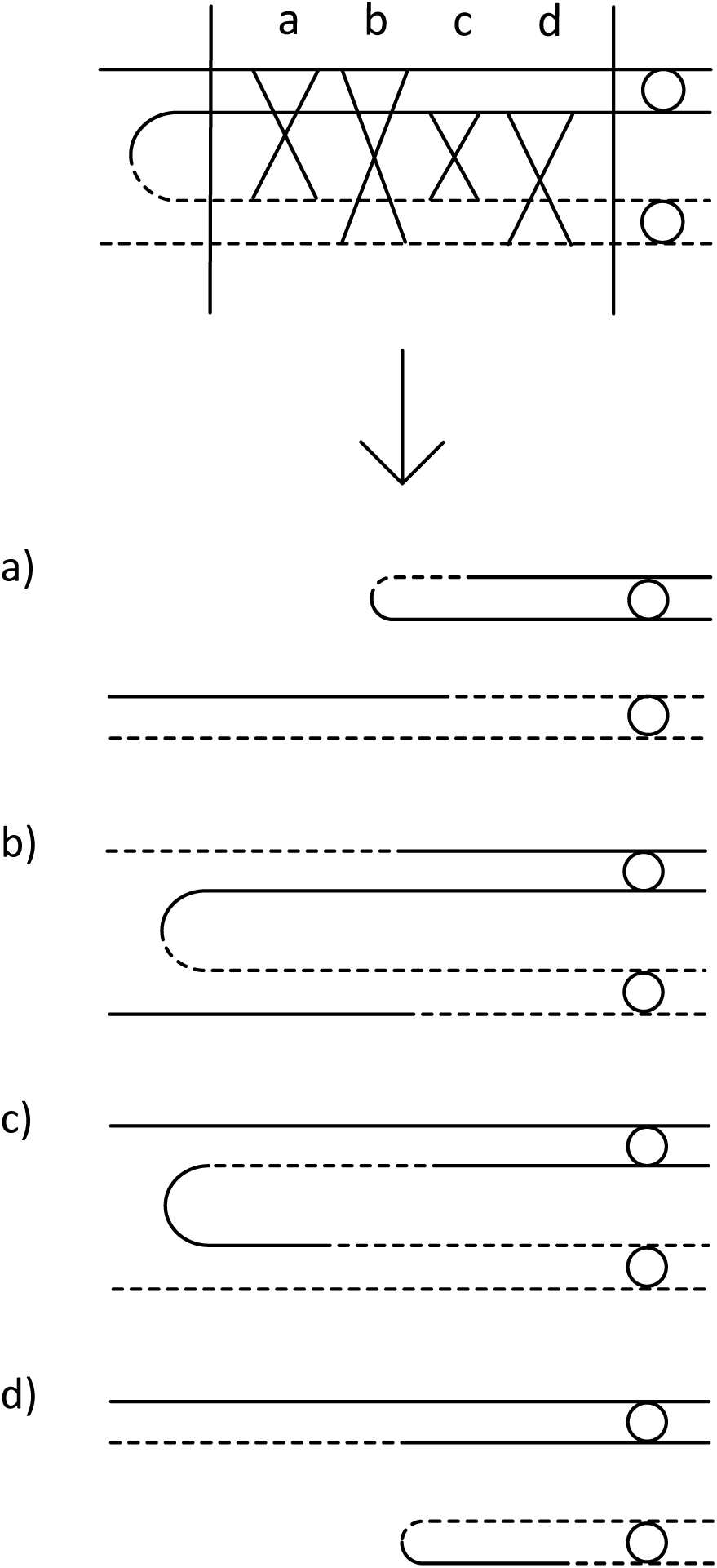
The effect of a single additional chiasma in the proximal region when the tetrad is originally in the configuration single anaphase I bridge. The figure shows that Pr{Z(x_1_, x_2_ + 1) = 1|Z(x_1_, x_2_) = 1} = 1/2 and Pr{Z(x_1_, x_2_ + 1) = 2|Z(x_1_, x_2_) = 1} = 1/2, which give the second row (zero-indexed row 1) of matrix **T**_𝕡_. The remaining rows are found in the same manner.

**Figure 9:**
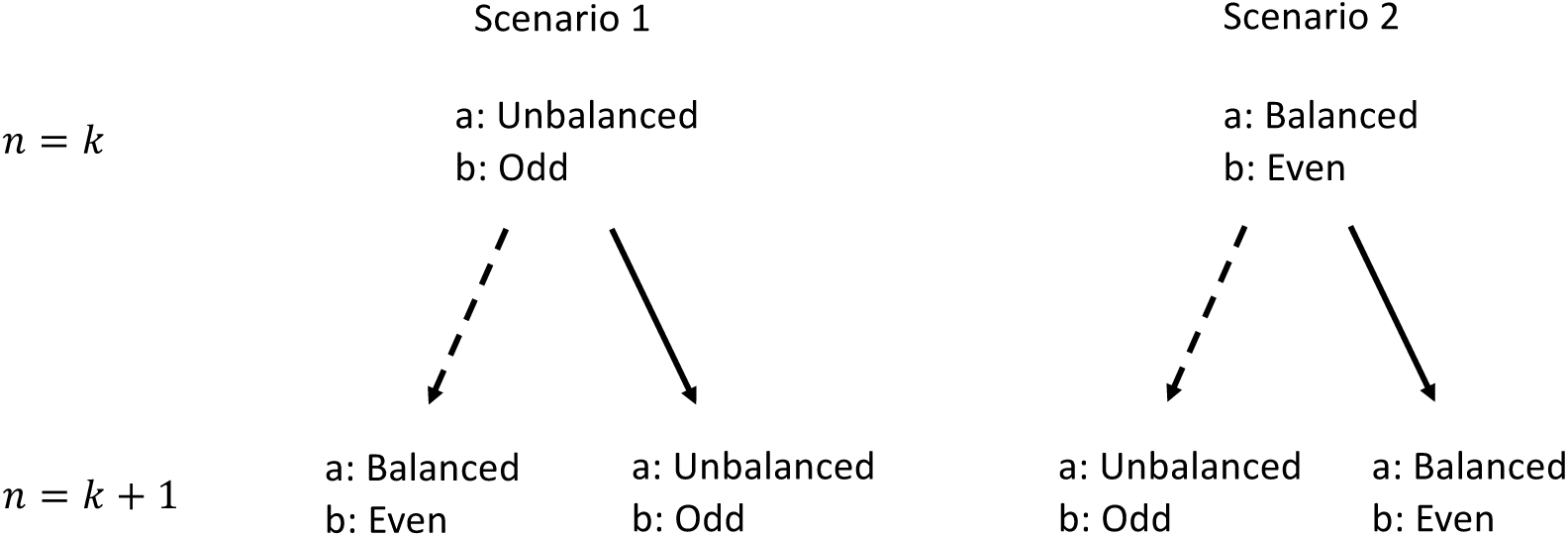
An induction proof of the statement (S) that a chromatid is unbalanced if and only if it shows recombination in an odd number of intervals within the inverted region. Keep in mind that an interval shows recombination if and only if it is involved in an odd number of chiasma events within the interval, and that a chromatid (in a heterokaryotype) is unbalanced if and only if it show recombination in the inverted region. Note first that the statement is trivially true when the inverted region consists of only a single interval (S_1_). Now assume for the sake of argument that the boundaries of the inverted region can be moved to include additional intervals, and that the statement is true when the inverted region consist of n = k intervals (S_k_). Hence, we must consider two scenarios: either the chromatid is unbalanced (involved in an odd number of chiasma events within the inverted region), which, the induction hypothesis states, must mean that it also shows recombination in an odd number of intervals within the inverted region (scenario 1), or the chromatid is balanced (involved in an even number of chiasma events within the inverted region), meaning that it show recombination in an even number of intervals within the inverted region (scenario 2). The figure shows the effect of including an additional recombinant (dashed arrow) or non-recombinant (solid arrow) interval in both of these scenarios (a: unbalanced/balanced chromatid, b: odd/even number of recombinant interval in the inverted region). Note that the statement is true for n = k + 1. Hence, since S_1_ is true and S_k_ implies S_k+1_, the statement must be true for all n > 0.

When all chromatids have an equal chance of becoming gametes, the sterility and recombination pattern probabilities for paracentric inversion heterokaryotypes is the same as for pericentric inversions. I will therefore refer to these cases collectively as *standard inversions*. In the *linear meiosis* of females of *Drosophila* (Sturtevant and Beadle 1936, Roberts 1976) and *Sciara* (Carson 1946), the two unbalanced chromatids in an anaphase I tetrad are retained in the polar bodies, so the remaining two balanced chromatids are the only ones that can become gametes. Since a single chiasma event in the inverted region always generates an anaphase I tetrad (figure 6), this means that additional chiasma events in either the inverted or proximal region are required to produce unbalanced gametes (figures 7 and 8). Hence, the sterility is significantly reduced. Furthermore, the set of recombination patterns that end up in an anaphase I tetrad is not a representative sample of the full set, so the gamete proportions are also affected. In the following, I will refer to the case of paracentric inversion heterokaryotypes with linear meiosis as *paracentric linear inversions*.

### 1.2 Chiasma interference

A *renewal process* (see Ross 2014, chapter 7) is as a stochastic process that represents the number of events (of a given type) that occur in an interval of a given length, when the distances between neighboring events are all drawn from the same *interarrival distance distribution*. I will say that the events *occur independently* or *are independent* if the probability of observing a given number of events in any given interval is independent of the number of events in all other disjoint intervals. The renewal process that possesses this attribute is known as the *Poisson process* (Ross 2014), in which the interarrival distance distribution is exponential, and the number of events in an interval of a given (genetic) length follows a Poisson distribution, so that

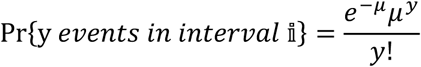

where *μ* is the expected number of events in interval 𝕚. If the chiasma events occur independently, they are distributed according to a Poisson process, and we say that there is *no chiasma interference*. This seems to be the case in some organisms, e.g. *Schizosaccharomyces pombe* (Munz 1994), but these are exceptional (see Berchowitz and Copenhaver 2010 for a review). I will refer to the model that assume no chiasma interference as *the Poisson interference model*, as first described in Haldane (1919). If chiasma events do not occur independently – i.e. if the probability of observing a given number of chiasma events in one interval depends on the number of chiasma events in a disjoint interval – we say that there is *chiasma interference*. A common measure of the degree of interference is the *coefficient of coincidence* (Muller 1916, Foss et al. 1993), one version of which is defined as

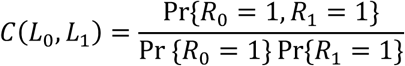

where 𝕚_0_ and 𝕚_1_ are two adjoining intervals of genetic length *L*_0_ and *L*_1_ (This is equivalent to *S*_*3*_ in Foss et al. 1993, except that they impose the additional constraint that *L*_0_ = *L*_1_). Note that if the chiasma events occur independently, then Pr{*R*_0_ = 1, *R*_1_ = 1} = Pr{*R*_0_ = 1} Pr{*R*_1_ = 1} and *C*(*L*_0_, *L*_1_) = 1 for all values of *L*_0_ and *L*_1_. It is common to distinguish between *positive chiasma interference*, in which a chiasma in one location impede (i.e. makes less likely) the generation of a chiasma in a nearby location (*C*(*L*_0_, *L*_1_)<*1* for small *L*_0_, *L*_1_), and *negative chiasma interference*, in which a chiasma in one location facilitates (i.e. makes more likely) the generation of a chiasma in a nearby location (*C*(*L*_0_, *L*_1_)>*1* for small *L*_0_, *L*_1_). In practice, chiasma interference is almost always positive and gradually decreasing with distance, i.e. *C*(*L*_0_, *L*_1_) is typically close to zero for small *L*_0_, *L*_1_ and approaches 1 as *L*_0_, *L*_1_ are increased (Berchowitz and Copenhaver 2010, Foss et al. 1993).

The existence of chiasma interference implies that the information that a chiasma has occurred at one location on a chromosome must somehow propagate to nearby locations where it interferes with (i.e. influences the probability of) the generation of other chiasmata. I will refer to this information as *the interference signal*. Exactly how the interference signal manifests itself physically is poorly understood (Hillers 2004, Berchowitz and Copenhaver 2010); suggestions include a hypothetical polymer that grows out from each chiasma (King and Mortimer 1990), and the build-up and release (at chiasma locations) of physical stress along the chromosome (Kleckner et al. 2004, Wang et al. 2015). Foss et al. (1993) suggests a model, henceforth referred to as the *pure counting model*, where *intermediate events* occur according to a Poisson process (i.e. independently) with rate *2(m*+*1)* along the tetrad, but each such event is subsequently resolved as either a chiasma event – with both crossing over and gene conversion – or what I will refer to as a *dummy event* – with gene conversion but not crossing over – in a strict sequence so that there are always *m* dummy events between each pair of chiasma events. Letting *C* denote an intermediate event, *C*_*x*_ a chiasma event and *C*_*0*_ a dummy event, the counting model hence postulates that for, say, *m = 2* the sequence …*C*_*x*_*C*_*0*_*C*_*0*_*C*_*x*_*C*_*0*_*C*_*0*_*C*_*x*_*C*_*0*_*C*_*0*_… with exponential interarrival distances between the *C’s* will be repeated along the span of the region of interest. This implies a chiasma interarrival distance distribution equal to the sum of *m*+*1* exponential distributions (m dummy events plus one chiasma event); in the literature on chiasma interference, this distribution is variously referred to as the chi-squared distribution (e.g. Zhao et al. 1995a) or the (scaled) Erlang distribution (Nolan 2017). Interference in the pure counting model depends on genetic, as opposed to physical, distance and is always positive (Lange et al. 1997) and stronger for higher *m;* the latter two attributes follow from the central limit theorem and the law of large numbers, as illustrated in figure 2. Note that the chiasma events occur independently if *m = 0*, because then all (independent) intermediate events are resolved as chiasma events. The Poisson interference model, in which there is no chiasma interference, can therefore be thought of as a special case of the counting model.

There are two ways of interpreting the pure counting model; either as a literal description of the physical manifestation of the interference signal – a “machine that can count”, in the words of Foss and Stahl (1995) – or as a mathematical abstraction that, disregarding gene conversions, models the distribution of chiasma events without making any claims as to how interference actually works. The latter interpretation is foreshadowed in the mathematically equivalent models of Cobbs (1978) and Stam (1979), among others (see McPeek and Speed 1995 for a brief historical overview). Foss et al. (1993), however, clearly favor the former interpretation, and they support this view by presenting a remarkably good fit between the predicted and observed coefficients of coincidence for *Drosophila* and *Neurospora* when *m*, crucially, is independently estimated from the ratio of the number of gene conversions to the number of chiasmata in a different dataset. Additional evidence for the accuracy of the pure counting model’s prediction of the distribution of chiasma events in at least some species is given in Lande and Stahl (1993), Zhao et al. (1995), and McPeek and Speed (1995). One prediction of the literal “machine that can count” interpretation – that a region enclosed by two chiasma events will be enriched for gene conversions without crossing over – is, however, not fulfilled in *Saccharomyces cerevisiae* (Foss and Stahl 1995). Stahl et al. (2004) and Malkova et al. (2004) suggest that this discrepancy is due to the presence of two distinct and independent types of chiasmata, with and without interference, in *S. cerevisiae* and some (but not all) other organisms, and they provide evidence to that effect (see also Berchowitz and Copenhaver 2010). I will refer to the chiasma interference model that incorporates the none-interference type in addition to the interference (counting) type of chiasma events as the *two-pathway counting model*. This model has proved a significantly better fit to data from e.g. *Arabidopsis thaliana* (Copenhaver et al. 2002) and humans (Housworth and Stahl 2003), compared to the pure counting model. Another suggested generalization, anticipated in Foss et al. (1993), is the *Poisson-skip model* of Lange et al. (1997), in which the number of dummy events between each chiasma event is drawn from a probability distribution. Among the advantages of this model is that allows for more fine-tuned modelling of interference strengths, and that it allows for negative as well as positive chiasma interference. For clarity, I will henceforth refer to the Poisson-skip model as the *stochastic counting model*.

In keeping with the deficiency of evidence for any of the proposed theories (Hillers 2004, Berchowitz and Copenhaver 2010), I will in this study remain agnostic about the physical manifestation of the interference signal and, disregarding gene conversions altogether, treat the dummy events as useful mathematical abstraction that may or may not actually correspond to sites of gene conversion without crossing over. On that basis, I will here suggest a further generalization of the four models introduced in this section – the *Poisson model, the pure counting model, the two-pathway counting model* and the *stochastic counting model* – which I will refer to as the *general counting model* or just *the general model* for short. I will then use this model to derive expressions for the coefficient of coincidence (theorem 1), recombination pattern probabilities in homokaryotypes (theorem 2), a closed-form version of theorem 1 in a special case (theorem 3), the sterility of standard inversion heterokaryotypes (theorem 4), the recombination pattern probabilities in standard inversion heterokaryotypes (theorem 5), the sterility of paracentric linear inversion heterokaryotypes (theorem 6) and the recombination pattern probabilities of paracentric linear inversion heterokaryotypes (theorem 7). Note that the expressions give recombination pattern probabilities of balanced gametes as the proportion among all gametes and not (as in Navarro et al. 1997) as a proportion of balanced gametes only; to get the latter, simply divide by 1 − ζ, where ζ is the sterility calculated by using the relevant theorem.

## 2 The model

As in the two-pathway counting model, I postulate two independent chiasma-generating pathways. The *type I pathway* is without chiasma interference, and generate only *type I chiasma events* according to a Poisson process. The *type II pathway* is (potentially) with interference, and generate *type II intermediate events* according to a Poisson process; these are subsequently resolved as either *type II chiasma events* or *type II dummy events*, where the number of the latter following each instance of the former is drawn from a user-defined probability distribution. That is, the type II chiasma events occur according to the stochastic counting model. It will be convenient to give the different events symbols for easier reference, in particular when these have to be incorporated into mathematical expression. In the following I will denote the type I chiasma events, type II chiasma events, type II dummy events, and type II intermediate events as *X*′, *X*″, *O*″, and *C*″, respectively (note that the *C*″ events comprises the union of all *X*″ and *O*″ events.). The union of all chiasma events of either type will be denoted *X*, and the union of all events – the *intermediate events* (without the *type II* qualifier) – will be denoted *C*. The same symbols with a subscript indicating an interval will serve as random variables representing the number of the event in question within that interval. Hence, I can, for example, write “the probability of observing one or more type I chiasma events in interval 𝕚 is *p*” as 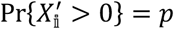, or “the expected number of type II intermediate events in interval 𝕚 is *λ*” as 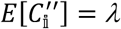.

In the homokaryotype case, the general model can be described by three user-defined sets of parameters. ***μ*** = {*μ*_0_, *μ*_1_, *μ*_2_, …, *μ*_𝔫−1_} and ***λ*** = {*λ*_0_, *λ*_1_, *λ*_2_, …, *λ*_𝔫−1_} are the expected number of type I chiasma events and type II intermediate events, respectively, in each of the 𝔫 intervals, so that 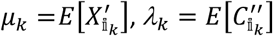 *for k* = 0,1,2 … 𝔫 − 1. Keep in mind that the 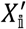 and 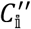 events are both Poisson distributed. The third set of parameters is the distribution of probabilities for observing a given number of consecutive *O*″ events following a *X*″ event. I will call this distribution the *intervening O*″*events distribution*, and denote it ***γ*** = {*γ*_0_, *γ*_1_, *γ*_2_ … *γ*_*m*_}, so that *γ*_*q*_ (*for q* = 0,1,2 … *m*, 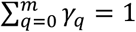) gives the probability of observing *q* consecutive *O*″ events following a *X*″ events, or, equivalently, *q* intervening *O*″ events between a pair of *X*″ events. One *cycle* hence consists of one *X*″ event and *q O*″ events, where *q* is drawn from ***γ***, so that the expected total number of *C*″ events in a cycle is 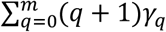. I will assume that there is a user-defined finite upper limit, *m*, on the possible number of intervening *O*″ events, so that *γ*_*q*_ = 0 *for q* > *m*. This implies that the genetic length of an interval 𝕚_*k*_ is given by

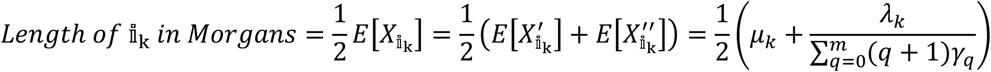

Note that the general counting model reduces to the stochastic counting model in the special case where *μ*_*k*_ = 0 *for k* = 0,1,2 … 𝔫 − 1, i.e. when there are no type I chiasma events; and to the two-pathway counting model in the special case where *γ*_*m*_ = 1, *γ*_*q*_ = 0 *for q* ≠ *m*, i.e. when there are strictly *m* intervening *O*″ events between all consecutive pairs of *X*″ events. Figure 2 presents the general counting model, the two-pathway counting model, the stochastic counting model, the pure counting model, and the Poisson model as a model hierarchy where each downwards arrow pointing from model *a* to model *b* indicate which parameters you have to restrict (and how) in model *a* to get model *b* as a special case. I will in this text sometimes refer to all these collectively as *the counting models*, and use the term *pure counting model* when referring specifically to the model suggested by Foss et al. (1993).

During meiosis for chromosomes heterozygous for a chromosomal inversion, the inverted region can form a homosynaptic *inversion loop* (figure 4) inside and outside of which the formation of chiasmata is partly or fully inhibited (Novitski and Braver 1954, Coyne et al. 1991, 1993, Navarro and Ruiz 1997, Jaarola et al. 1998, Anton et al. 2005, Pegueroles et al. 2010, del Priore and Pigozzi 2015). I will henceforth define the *inhibition factor* for interval 𝕚_*k*_, denoted *d*_*k*_ ∈ ℝ_≥0_, so that if the expected number of *X*′ and *C*″ events in interval 𝕚_*k*_ is *μ*_*k*_ and *λ*_*k*_, respectively, in a homokaryotype, then the corresponding values in a heterokaryotype are *d*_*k*_*μ*_*k*_ and *d*_*k*_*λ*_*k*_. The two boundaries of the inverted region, the *breakpoint boundaries*, can for our purpose be thought of as loci that serve as interval boundaries in the same way that other loci do, so that e.g. the interval between the left breakpoint boundary and the leftmost loci within the inverted region (or the right breakpoint boundary if there are no loci within the inverted region) is an interval in the same respect that the intervals enclosed by two non-breakpoint loci are. Since inversions affect the rate of chiasma generation also in interval outside of the inverted region (e.g. Pegueroles et al. 2010), *d*_*k*_ also applies for such intervals. I will adopt the convention that *E*[] always refer to expectation in a homokaryotype, so that e.g. 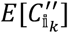 and 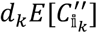 gives the expected number of type II intermediate events in interval *k* in a homokaryoype and heterokaryotype, respectively. This notation allows different intervals within and outside of the inverted region to be experience inhibition to different degrees, depending, for example, on their distance from the breakpoints.

For pericentric inversion heterokaryotypes, I will divide the region of interest into three subregions for easier reference. *The left region* is the region comprising all intervals to the left of the inverted region, *the inverted region* is, as before, the region comprising all intervals captured by the inversion, and *the right region* is the region comprising all intervals to the right of the inverted region. These three regions will be denoted 𝕕, 𝕙 and 𝕣, respectively, so that e.g. 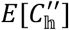 and 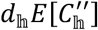 denotes the expected number of type II intermediate events in the inverted region as a whole in a homokaryotype and heterokaryotype, respectively. The number of intervals in 𝕕, 𝕙, and 𝕣 will be denoted 𝔡, 𝔥, and 𝔯, so that the total number of intervals is 𝔫 = 𝔡 + 𝔥 + 𝔯. As before, 𝕚_*k*_ refer to zero-indexed interval number *k* from the left (one-indexed interval *k* + *1*), so from the preceding we can unambiguously deduce that 𝕚_*k*_, *k* = 0,1,2 … 𝔡 − 1 is an interval in the left region, and 𝕚_*k*_, *k* = 𝔡, 𝔡 + 1, 𝔡 + 2 … 𝔡 + 𝔥 − 1 is an interval in the inverted region, and 𝕚_*k*_, *k* = 𝔡 + 𝔥, 𝔡 + 𝔥 + 1, 𝔡 + 𝔥 + 2 … 𝔫 − 1 is an interval in the right region. For easier reading, I will when convenient adopt the convention that an 𝕕, 𝕙, and 𝕣 with subscript index *k* refer to the same interval as 𝕚_*k*_, the only difference being the additional explicit (and redundant) information about which of the three subregions the interval belongs to. For example, 𝕚_*k*_ and 𝕙_*k*_ refer to the same interval, zero-indexed number *k* from the left in the region of interest, but in the latter case you can immediately see that the interval is in the inverted region without having to scrutinize the index.

For paracentric linear inversion heterokaryotpes, I will divide the region of interest into four subregions from left to right: *the distal region* comprises all intervals to the left of the inverted region, *the inverted region* is as before, the *proximal region* comprises all intervals between the inverted region and the centromere (which, for our purpose, serve as a loci), and the *right region* comprises all intervals to the right of the proximal region. These will in the following be denoted 𝕕, 𝕙, 𝕡, and 𝕣, respectively, with the number of intervals in each denoted 𝔡, 𝔥, 𝔭 and 𝔯. Otherwise the notation is the same as for pericentric inversions.

Since the order of the intervals within the inverted region is inverted in one homologue with respect to the other, there are at least two equivalent paths through which the interference signal can travel across this region (figure 5). This raises the empirical problem of how the interference signal travels through inversion breakpoint boundaries, which to my knowledge is not addressed in the literature – unsurprisingly, given that the physical manifestation of the interference signal is yet to be satisfactory described (Hillers 2004, Berchowitz and Copenhaver 2010). In my model, I suggest either to calculate the recombination pattern probabilities under the assumption that the interference signal travels through either of the two paths with equal probability, or to do so under the assumption that the signal is blocked by the inversion boundaries so that the number of chiasma events inside the inverted region is independent of the number outside of this region (there is evidence that this occurs in at least some cases; Gorlov and Borodin 1995). The Boolean parameter *α* can be set to 1 or 0 to facilitate either of these possibilities. Note that in the former case I have for notational simplicity only included one of the two possible paths in the theorems; to get the desired result, do the calculations twice with the order of the intervals inside the inverted region reversed and average the results. When the interference signal is blocked by the inversion boundaries (*α* = 0), the directionality of the inverted region is irrelevant under the assumption of stationarity (see the next section), so only one calculation needs to be performed.

### 2.1 Phases and stationarity in the general model

In the following, I will define the *phase* at any location on a tetrad as the number of *O*″ events between that location and the nearest *X*″ event to the *right*. I will furthermore define 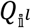 and 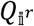 as random variables representing the phase at the left and right boundary, respectively, of interval 𝕚, and more generally *Q*_*a*_ as a random variable representing the phase at a location *a*. A *phase distribution* is now a probability distribution that gives the probabilities of observing the individual possible phases at a given location. The *stationary phase distribution* is the phase distribution satisfy the equation set

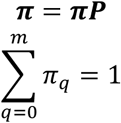

where ***π*** = (*π*_0_ π_1_ … π_m_) = (Pr{*Q*_*a*_ = 0} Pr{*Q*_*a*_ = 1} … Pr{*Q*_*a*_ = *m*}) and ***P*** is the Markovian transition matrix that transforms the phase distribution at a given location *a* into the phase distribution at a location *b*, when there is one *C*″ event between *a* and *b*, i.e.

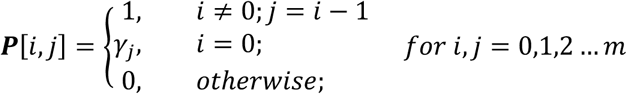

(See the appendix for a note on the matrix notation used in this text). Solving this equation set gives

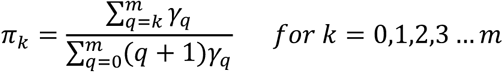

which is the stationary phase distribution for the general model. This distribution is important because if it is given that the phase distribution at any given location *a* corresponds to this distribution, then the phase distribution at all other locations *b* also corresponds to this distribution. This means that if we *assume stationarity*, then all regions that are described by the same set of **λ, σ** and **γ** values are fungible, regardless of their position on the tetrad. It also means the *left* and *right* boundaries of any interval or region have the same phase distribution, so that it does not matter which we call *left* and which we call *right*. In other words, reversing the numbering of the intervals in the region of interest does not change the results.

Cobbs’ (1978) and Stam’s (1979) early analyses of (the mathematical equivalent of) the pure counting model were complicated by the assumption of a non-stationary phase distribution (though they both considered the stationary phase distribution as a special case), which were motivated by Mather’s (1938) hypothesis that the process of chiasma generation starts at the centromere and proceed from there in both directions, implying that the phase at the centromere is non-stationary. This hypothesis was in turn motivated by early results indicating that the interference signal is blocked by the centromere, meaning that chiasma on one side of the centromere does not interfere with chiasma on the other. However, a more recent analysis by Colombo and Jones (1997) indicate that the results cited by Mather are merely statistical artefacts, and that, in contradiction to Mather’s predictions, interference does in fact work across the centromere in the same way as in other regions. This leaves it unclear exactly how and where the process of chiasma generation starts; or as Colombo and Jones (1997, p. 226) put it, “If chiasma formations does not start from the centromere, or end against the centromere, where does it start from or end? Or, to put it bluntly, does it start from anywhere?” Given that we do not know the answers to these questions (Hillers 2004, Berchowitz and Copenhaver 2010), it seems to me that assuming a stationary phase distribution, as more recent treatments of chiasma interference tend to do, is as plausible and parsimonious as any other alternative. I will therefore do so in the rest of this text. The predictions of a stationary and non-stationary model are in any case not very different (Lande and Stahl 1993).

## 3 Results

### 3.1 Theorem 1: The coefficient of coincidence for the general model

The coefficient of coincidence for the general model is given by

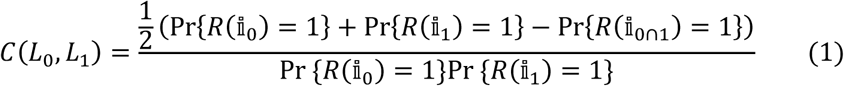

where 𝕚_0∩1_ is the interval between the left boundary of 𝕚_0_ and the right boundary of 𝕚_1_, and

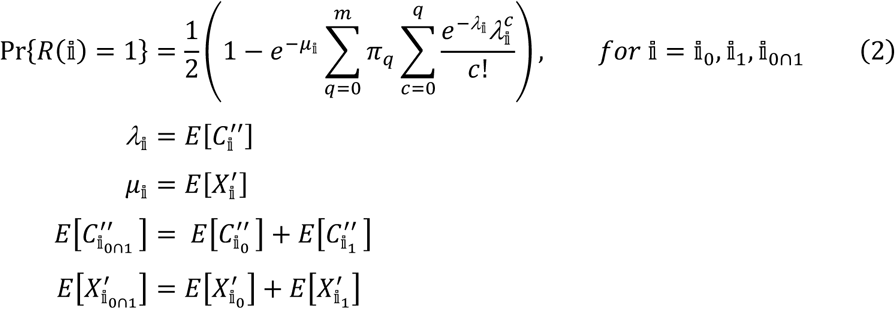

*Proof:*

Equation (1) follows from the definition of *C*(*L*_0_, *L*_1_) (section 1.2) and the relation

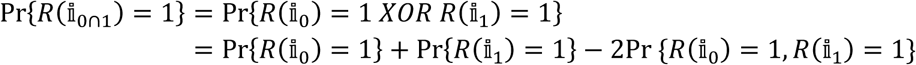

(Foss et al. 1993). To get equation (2), first note that

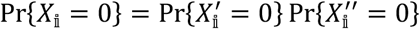

where

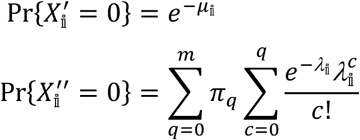

The latter expression sums over all possible phases *q* at the left boundary of 𝕚 weighted by their stationary probabilities (*π*_*q*_), and multiplies each term with the probability of observing between 0 and q *C*″ events in the interval, which by the definition of phase means all consecutive *C*″ events up to, but not including, the first *X*″ event. Equation (2) now follows from Mather’s (1938) equation,

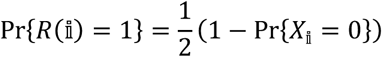

*QED*

### 3.2 Theorem 2: Recombination in homokaryotypes

Assuming stationarity and no chromatid interference, the probability of observing recombination pattern ***r*** on a homokaryotypic chromosome under the general counting model is given by

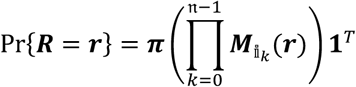

where

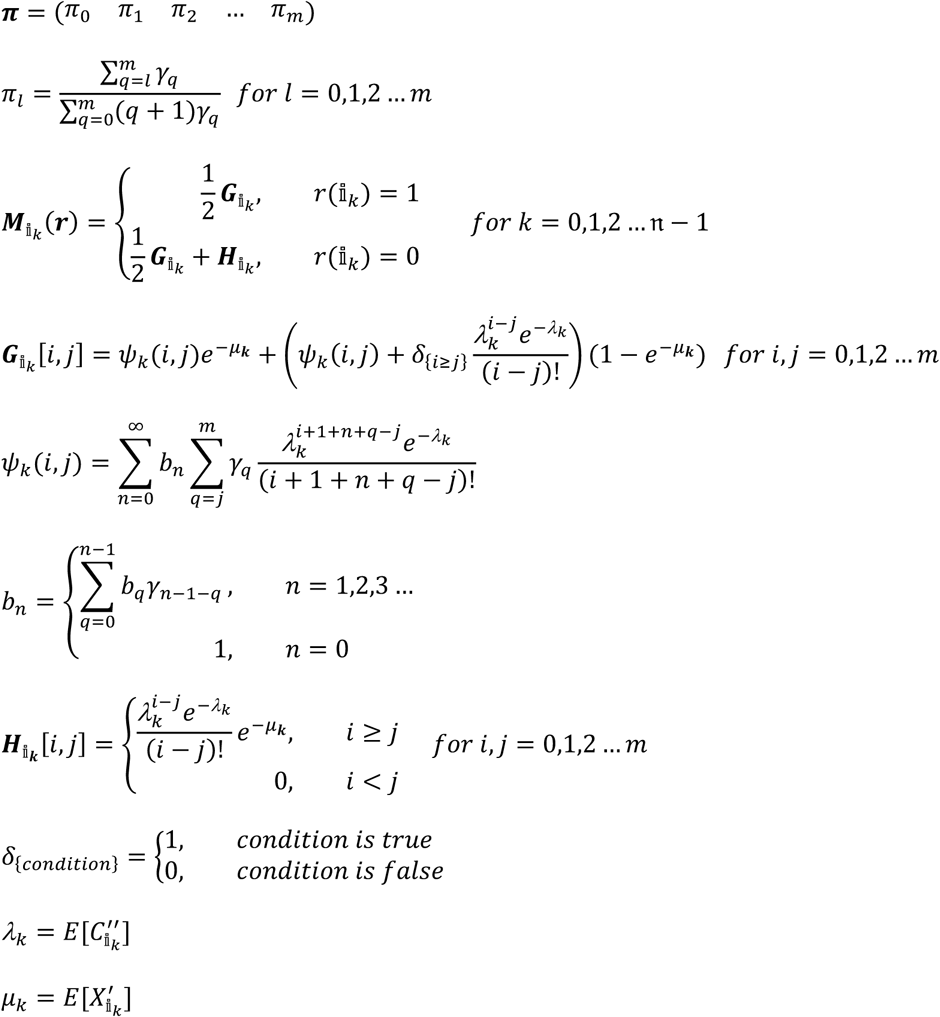

*Proof:*

As in Zhao et al. (1995), Lange et al. (1997) and Copenhaver et al. (2002), the key idea in theorem 1 is to sum over all possible phases at each boundary by means of matrix multiplications. The vector ***π*** gives the phase distribution at the leftmost boundary^2^, and element *i,j* of the 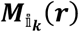 matrices gives the probability of observing the relevant recombination status (according to ***r***) *and* phase *j* at the right boundary of interval 𝕚_*k*_, *given* phase *i* at the left boundary of 𝕚_*k*_, i.e.

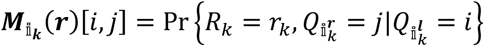

I will in this proof first show by induction that

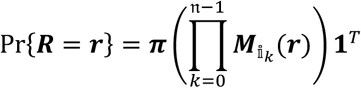

before I will find expressions for 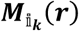 in terms of the model parameters.

For the induction proof, let *S*_*1*_ be the statement 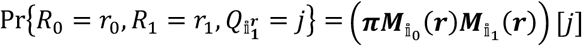, and the induction hypothesis *S*_*k-1*_ be

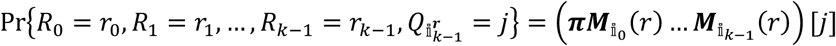

To prove *S*_*1*_, note first that

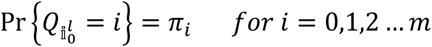

so

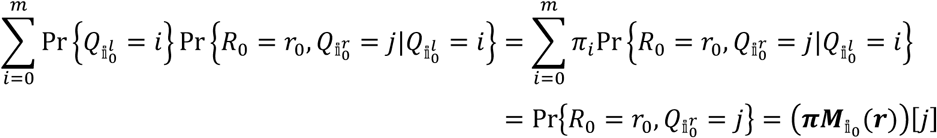

Using that 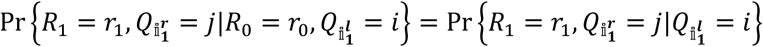 (because *R*_0_ is irrelevant if the phase at the left boundary of 𝕚_1_ is known) and that 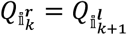 we have

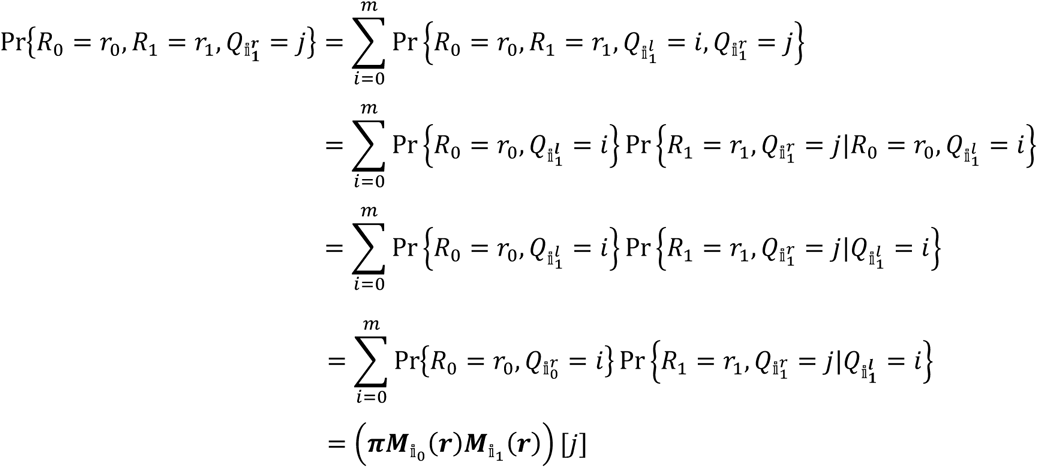

Hence, *S*_*1*_ is true. Now assume that the induction hypothesis *S*_*k-1*_ is true and observe that

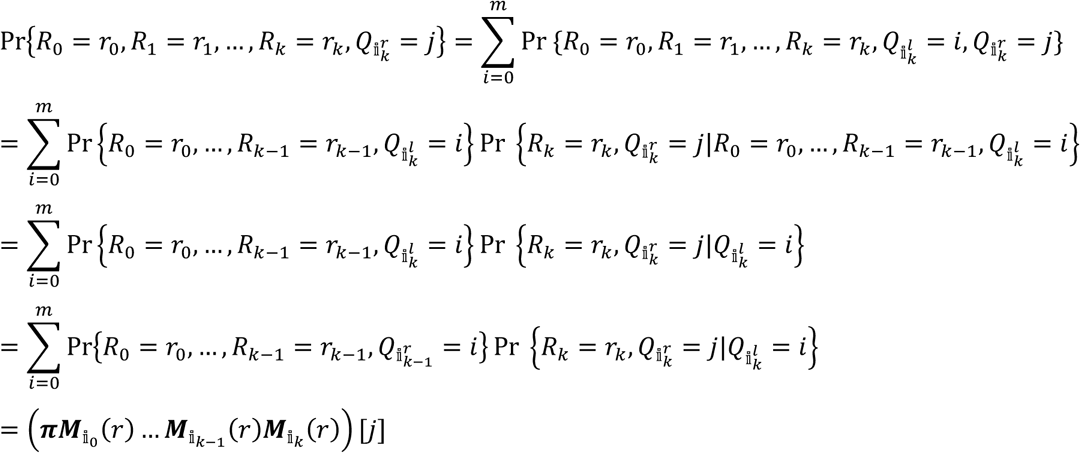

where the last equality follows from the induction hypothesis. Since *S*_1_ is true and *S*_*k*−1_ implies *S*_*k*_, is follows that *S*_𝔫−1_ must be true. Hence,

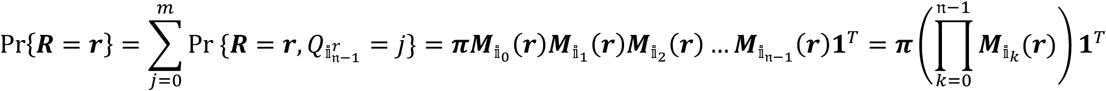

which concludes the induction proof.

The next step is to express the 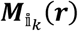 matrices in terms of the model parameters. I will do this by writing 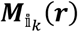 as a linear combination of the matrices 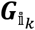 and 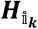 with elements

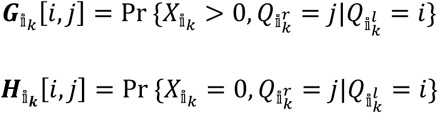

and then find such expressions for these matrices. For this end, I use Mather’s (1938) equation

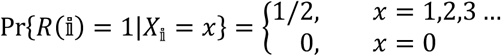

from which it follows that

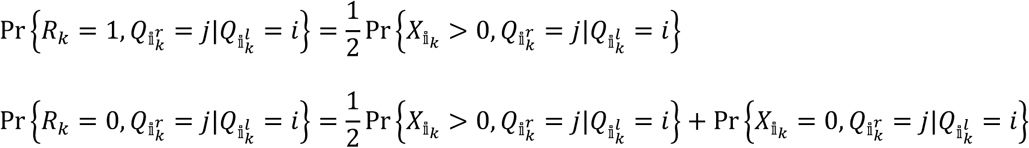

so

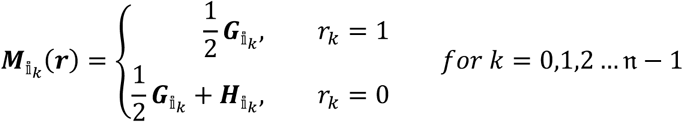

Element *i,j* of 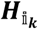 is equal to the probability of observing no chiasma events (of either type) in 𝕚_*k*_ *and* phase *j* at the right boundary, *given* phase *i* at the left boundary, which in the case *i* ≥ *j* is equivalent to the probability of observing 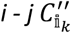 events multiplied by the probability of observing 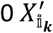 events. If *i* < *j*, it is impossible to arrive at phase *j* without at least one *X*″ event in the interval, so the probability is in this case 0. Hence,

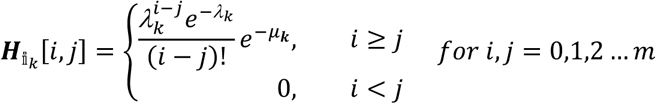

To find an expression for 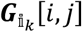, I first define the functions *ψ*_*k*_(*i, j*) as

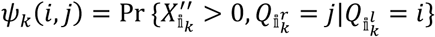

and *b*_*n*_ as

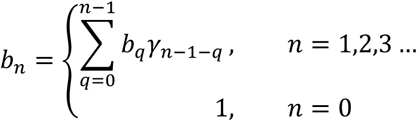

Note that *b*_*n*_, equivalent to *u*_*n*_ in Lange et al. (1997), gives the probability that the *n-th C*″event to the right of any *X*″event is also a *X*″event; as in Lange et al., it is derived by recursively conditioning on the index of the last *X*″ event before the *n*-th *C*″ event, with the base case *b*_0_ = 1. To get at least one 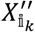 event and phase *j* at the right boundary, given phase *i* at the left boundary, one must observe in interval 𝕚_*k*_ (*i O*″ events to get to the first *X*″event) + (the first *X*″ event) + (*n* additional *C*″events of which the last is a *X*″ event) + (*q - j* additional *O*″events, where *q* is the number of *O*″events in the rightmost cycle of the interval, to end up in phase *j*), in total *i* + *1* + *n* + *q – j C*″ events. Hence, by summing over all possible values of *n* and *q*,

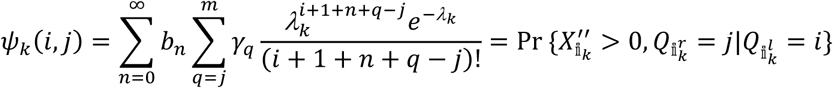

which is similar to the 1_{*j*>0}_ term in equation 6 in Lange et al. (1997) (though note that my ‘phase’ is defined differently from their ‘state’). By conditioning on the presence 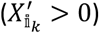 or absence 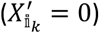 of *X*′ events in the interval,

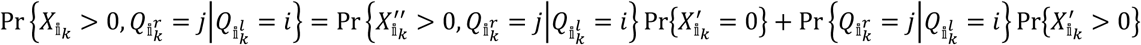

Since

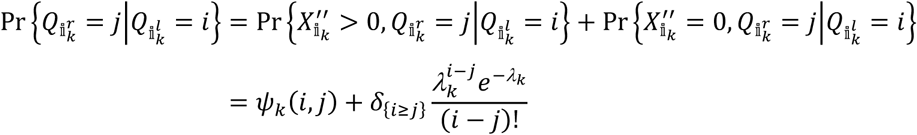

and

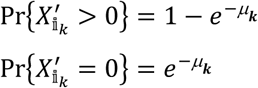

we now have that

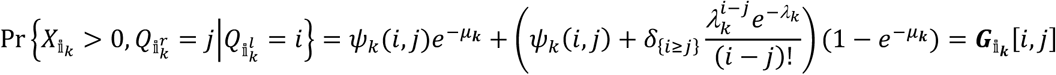

*QED*

### 3.3 Theorem 3: A closed-form version of the *G* matrices

#### 3.3.1 Prologue

Nolan (2017) derives a closed-form expression for the recombination pattern probabilities under the counting model. Building on the results from that paper, I will in this theorem do the same for the more general two-pathway counting model.

#### 3.3.2 Theorem

In the special case where *γ*_*m*_ = 1, *γ*_*q*_ = 0 *for q* ≠ *m* (i.e. the two-pathway counting model), the matrix 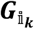 can be written in closed form as

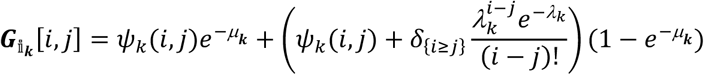

where

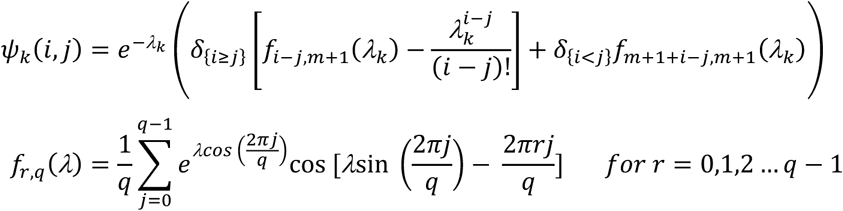

*Proof:*

The key to this theorem is the relation

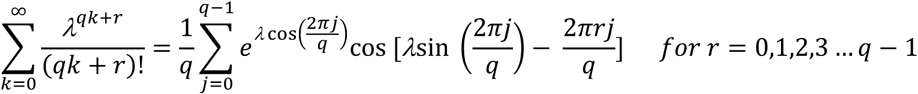

which is derived in Erdelyi (1955) and Nolan (2017). If *γ*_*m*_ = 1, *γ*_*q*_ = 0 *for q* ≠ *m*, then *ψ*_*k*_(*i, j*) reduces to

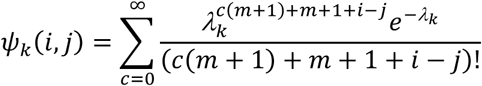

which we can now write as

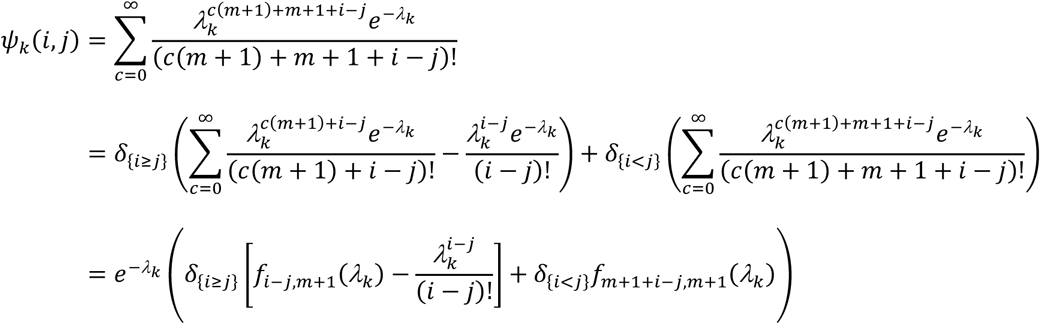

where

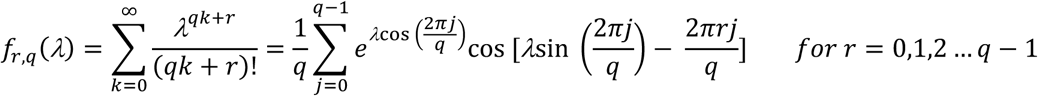

(note that *f* is not defined for *r ≥ q*)

*QED*

### 3.4 Theorem 4: The sterility of standard inversion heterokaryotypes

The sterility of an individual heterozygous for a standard inversion is given by

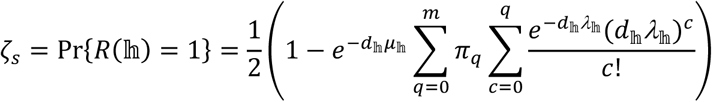

*Proof:*

For standard inversion heterokaryotypes, a chromatid is, as illustrated in figure 2.2, unbalanced if and only if it shows recombination in the inverted region. Since by assumption all chromatids, unbalanced or not, have an equal chance of becoming gametes, the sterility, ζ, of such a chromosome is simply given by ζ = Pr {*R*(𝕙) = 1}. The rest follows from the discussion in theorem 1.

*QED*

### 3.5 Theorem 5: Recombination in standard inversion heterokaryotypes

The probability of observing recombination pattern *r* on a chromosome with a standard inversion is given by

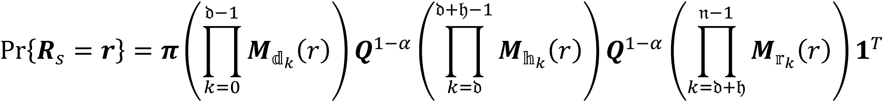

where

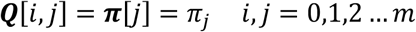

and the ***M*** matrices are the same as in theorem 1, except that the lambda and mu values are weighted by *d*_*k*_ so that e.g.

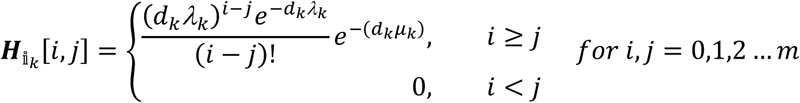

*Proof:*

Apart from the inhibition factors (*d*), the only difference between theorems 1 and 5 is the matrix ***Q*** that is inserted into the latter equation at the positions corresponding to the breakpoint boundaries. The parameter *α* indicate the presence (*α* = 1) or absence (*α* = 0) of chiasma interference across the breakpoint boundaries, so in the former case ***Q***^1−*α*^ = ***Q***^0^ = ***I***_*m*+1_ (i.e. the identity matrix). In the absence of interference across the breakpoint boundaries (*α* = 0), ***Q*** serves to redistribute the phase probabilities according to the stationary distribution (***π***). We can see why this is so by decomposing ***Q*** into the two matrices ***Q***′ and ***Q***″ so that

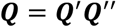

where

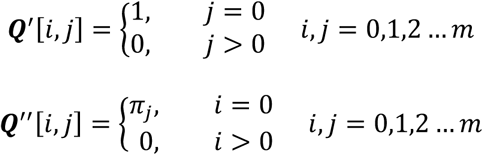

That is, ***Q***′ maps all phase probabilities (arbitrarily) to phase 0, whereas ***Q***″maps them from 0 to *j* in proportion to the stationary probabilities. This is equivalent to making the chiasma events in the three subregions mutually independent.

*QED*

### 3.6 Theorem 6: The sterility of paracentric linear inversion heterokaryotypes

#### 3.6.1 Prologue

Navarro et al. (1997) provide approximate expressions for the sterility and gamete proportions in paracentric linear inversion heterokaryotypes under the Poisson and pure counting interference models for a maximum of two loci in either the distal, inverted or proximal region. I will in this theorem and the next present exact infinite series expressions for the sterility and gamete proportions under the general interference model for an indefinite number of loci in each of the regions on the chromosome.

### 3.6.2 Theorem

The sterility of a paracentric inversion heterokaryotype with linear meiosis is given by

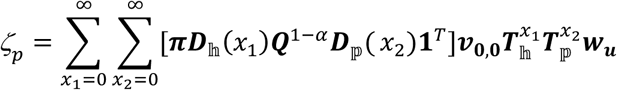

where

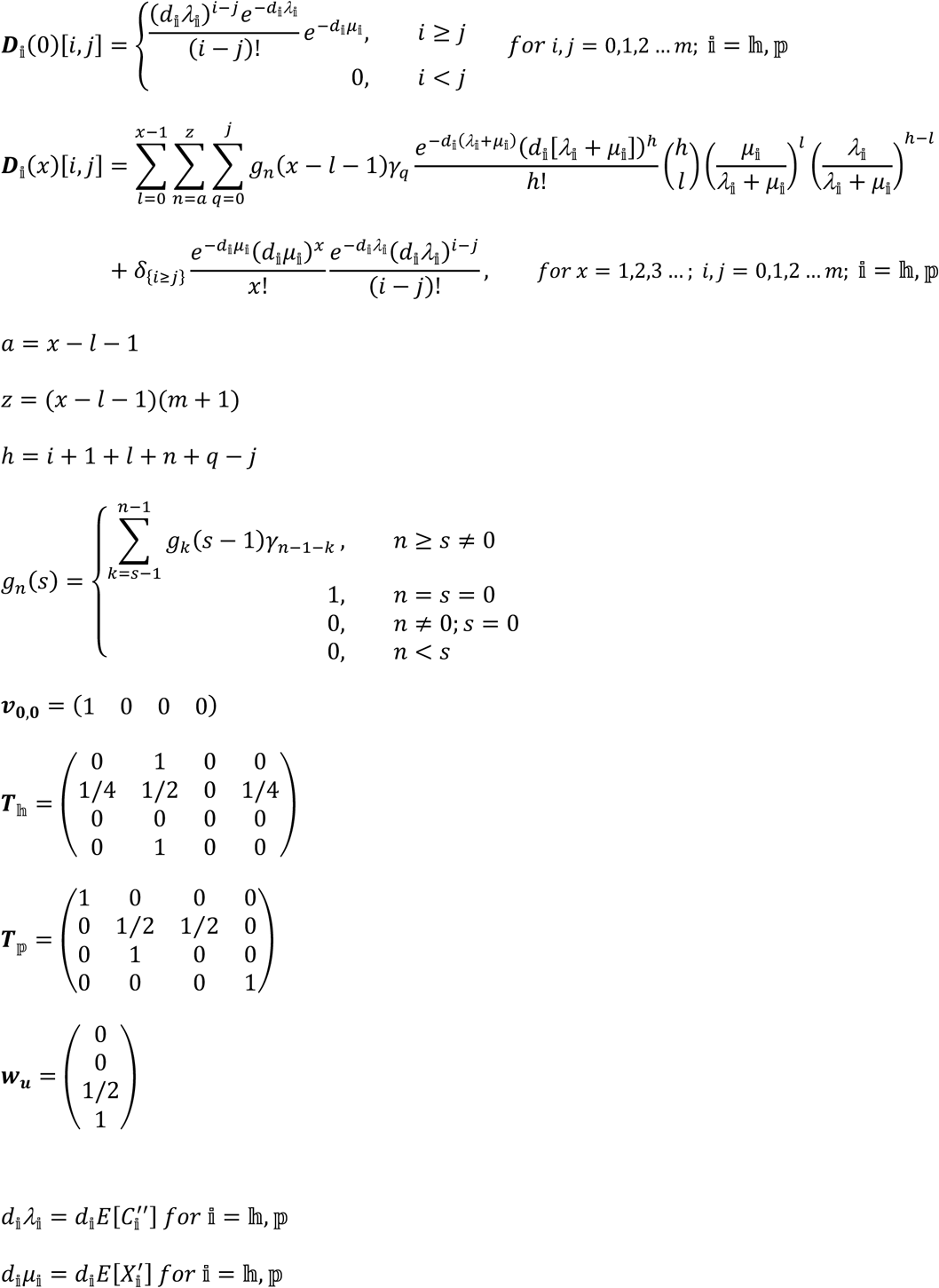

*Proof:*

The basic structure of this equation is

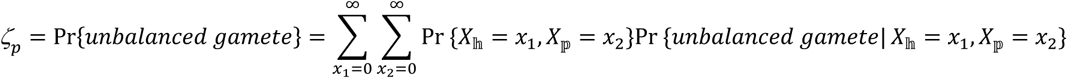

i.e. it finds the probability of an unbalanced gamete by conditioning on the number of chiasma events in each region. Pr{*X*_𝕙_ = *x*_1_, *X*_𝕡_ = *x*_2_} = ***πD***_𝕙_(*x*_1_)***Q***^1−*α*^***D***_𝕡_(*x*_2_)**1**^*T*^ is a special case of a more general expression that is shown in the proof for theorem 7. I therefore focus here on the conditional probability of observing an unbalanced gamete, Pr{*unbalanced gamete*|*X*_𝕙_ = *x*_1_, *X*_𝕡_ = *x*_2_}, which I will derive by means of a Markov chain, as follows. Consider a stochastic process 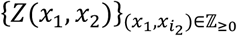 that represents the configuration of a tetrad when *X*_𝕙_ = *x*_1_, *X*_𝕡_ = *x*_2_ (i.e. when there are *x*_*1*_ chiasma events in the inverted region and *x*_*2*_ chiasma events in the proximal region), so that

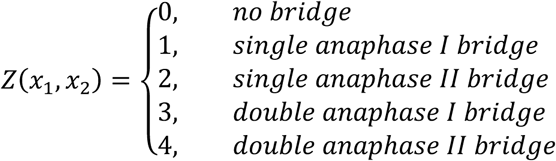

If I now define the matrices ***T***_𝕙_ and ***T***_𝕡_ so that

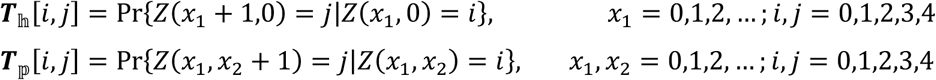

and the vector ***v***_0,0_ so that

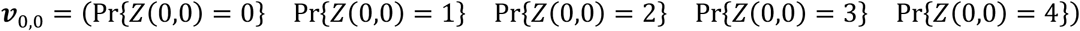

then the ***T***_𝕚_ matrices are the Markovian transition matrices whose element *i,j* give the probability of observing a tetrad in configuration *j* after considering an additional chiasma event at the right end of region 𝕚 (= 𝕙, 𝕡), given that the tetrad was in configuration *i* before considering that chiasma event and that there are no other chiasma event further to the right; and ***v***_**0**,**0**_ is the tetrad configuration distribution when there are no chiasma events in either interval. Accordingly,

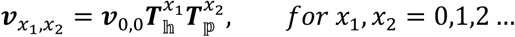

where

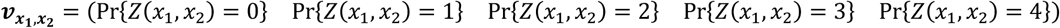

I now find Pr{*unbalanced gamete*|*X*_𝕙_ = *x*_1_, *X*_𝕡_ = *x*_2_} by weighting each configuration by its proportion of unbalanced gametes, i.e.

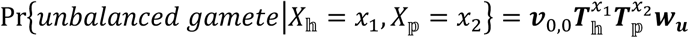

where

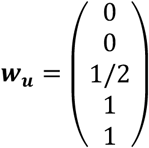

***T***_𝕙_ and ***T***_𝕡_ can be found by considering the resulting tetrads for the four possible combinations of non-sister strand involvements when an extra chiasma event is added at the right end of the appropriate interval of a tetrad that is originally in the given configuration (see figures 6 and 7). This results in

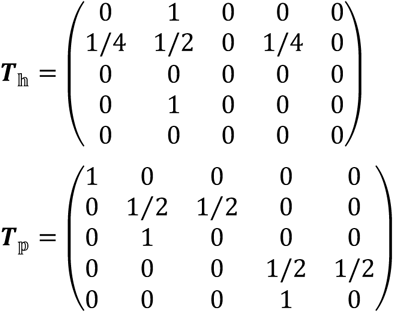

(Cobbs (1978) uses a similar line of reasoning to find the probability distribution of parental ditype, tetratype, and non-parental ditype tetrads given the number of chiasma events in a single colinear interval; note the similarity between his matrix ***T*** and my ***T***_𝕙_.) Since state 3 and 4 are identical for our purpose and form an absorbing class in ***T***_𝕡_, we can merge them into a single class *double bridge* to get the alternative (equivalent) expressions for ***v***_0,0_, ***T***_𝕙_, ***T***_𝕡_ and ***w***_***u***_ given in the theorem.

### 3.7 Theorem 7: Recombination in paracentric heterokaryotypes with linear meiosis

#### 3.7.1 Prologue

For this theorem we will need some additional terminology. I will say that a chromatid is in *state* 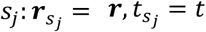 if it shows recombination pattern ***r*** *and* is in a tetrad with configuration *t*, and I will refer to 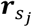 as the recombination pattern associated with state *s*_*j*_, and 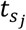 as the tetrad configuration associated with state *s*_*j*_. The *statespace* is hence the set of all possible combinations of recombination patterns and tetrad configuration, except that all states with unbalanced recombination patterns that also show recombination in any interval with index higher than 𝔡 + 𝔥 − 1 (i.e. the intervals in the proximal and rightmost regions) are excluded. As an example, with two intervals in the inverted region and one in the proximal region and zero intervals in the distal and rightmost regions (i.e. 𝔡 = 0, 𝔥 = 2, 𝔭 = 1, 𝔯 = 0), which means that there is one loci of interest in the inverted region and none in either of the other regions (note that this implies that 𝕚_2_ = 𝕡_2_ = 𝕡), we get the (arbitrarily numbered) statespace:

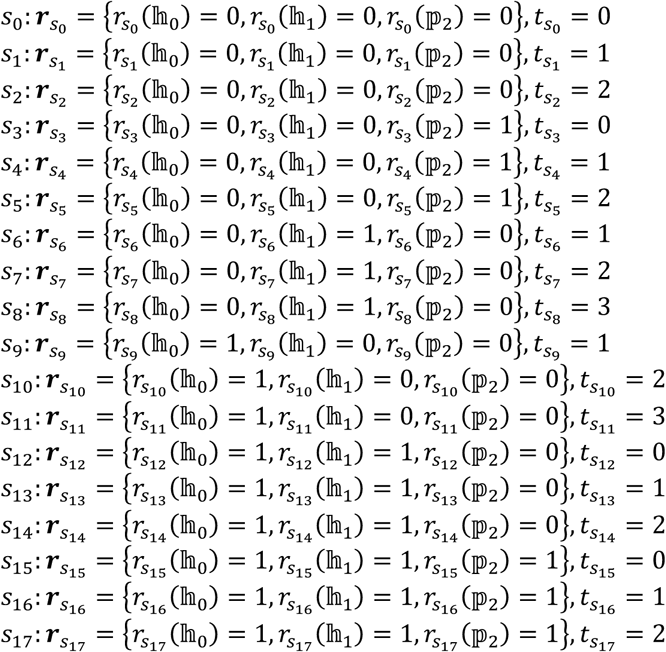

where

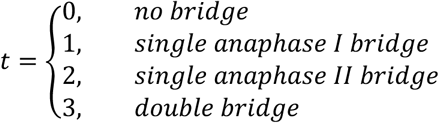

Note that there are no states for unbalanced patterns in *no bridge* tetrads or balanced patterns in *double bridge* tetrads; this is because if the tetrad is in the *no bridge* configuration, then none of the patterns associated with the tetrad can be unbalanced, and, similarly, if the tetrad is in the *double bridge* configuration, then none of the patterns associated with the tetrad can be balanced, so these combinations are impossible and hence not included in the statespace. The number of states – or, equivalently, the *size* of the statespace – will henceforth be denoted 𝔰; in the example above 𝔰 = 18.

### 3.7.2 Theorem

Let the statespace be as defined above. The probability of observing a gamete with recombination pattern ***r*** for a paracentric linear inversion heterokaryotype is now given by

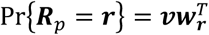

where

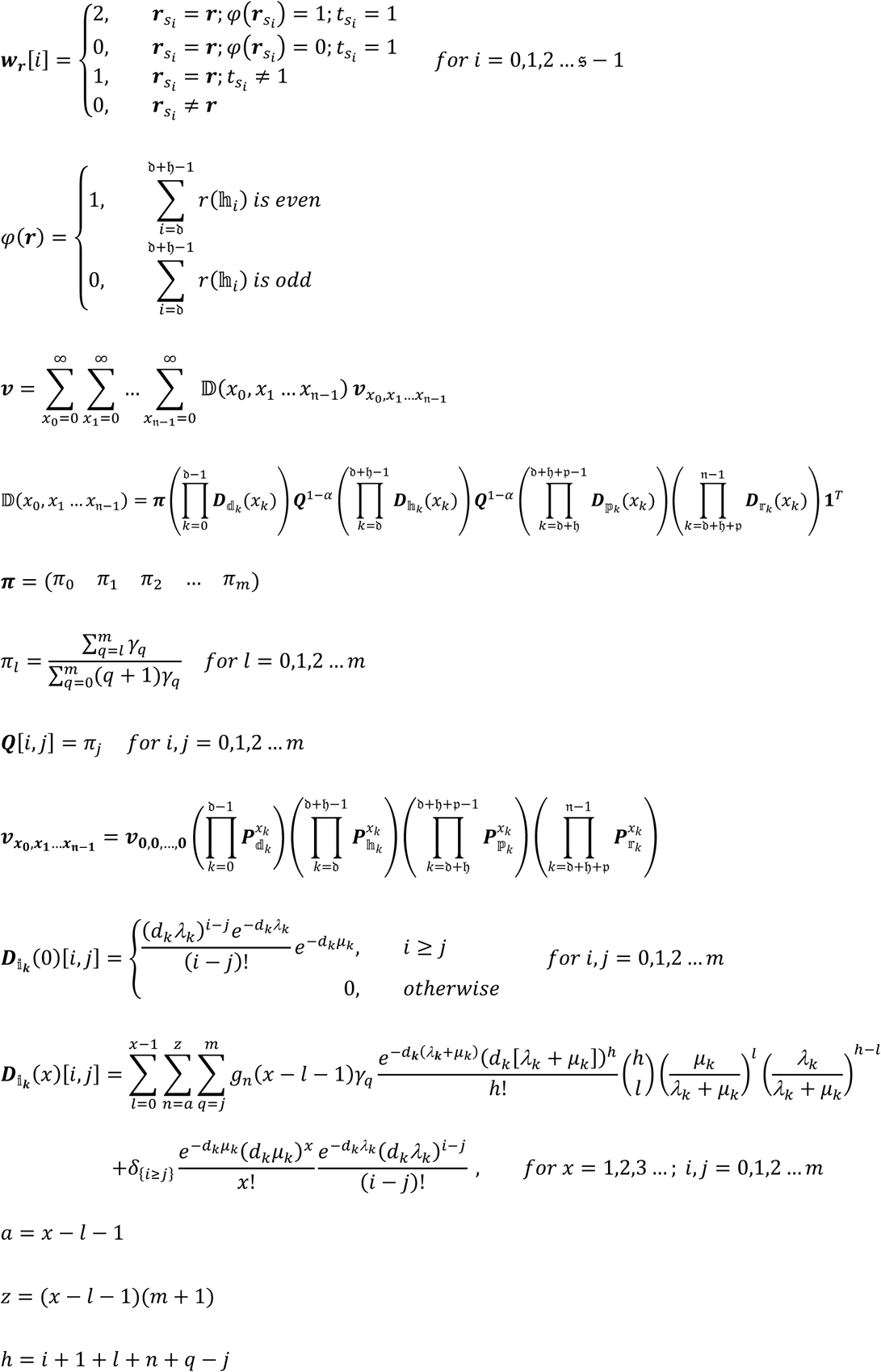

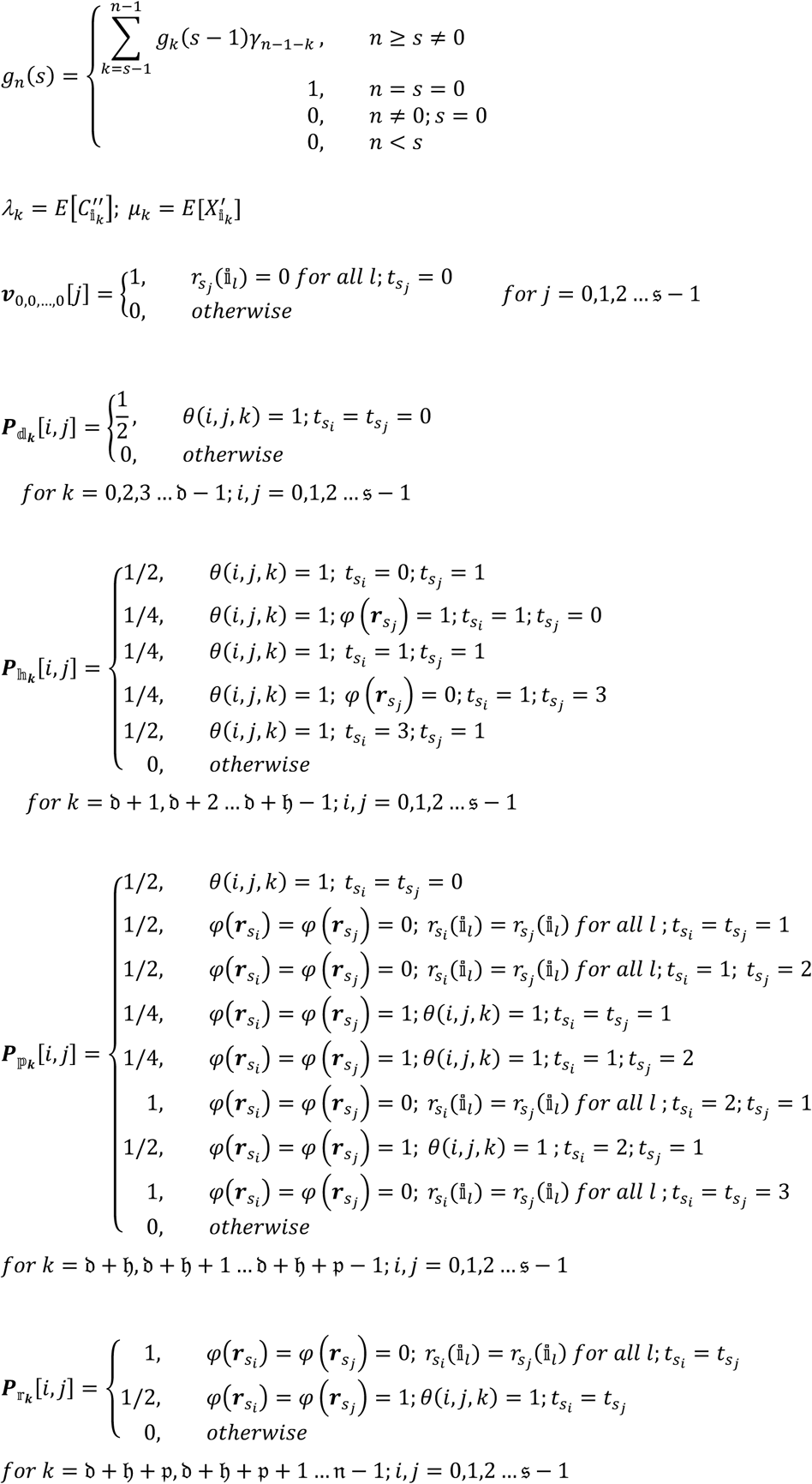

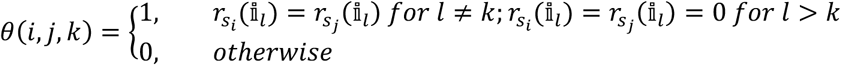

*Proof:*

Consider first a stochastic process 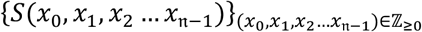 where *S*(*x*_0_, *x*_1_, *x*_2_ … *x*_𝔫−1_) represents the state of a chromatid (recombination pattern and tetrad configuration) when there are *x*_*0*_ chiasma events in 𝕚_0_, *x*_*1*_ chiasma events in 𝕚_1_, etc, so that

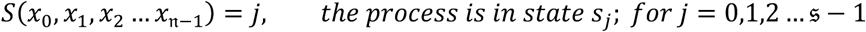

If we now define the vector ***v***_0,0,…,0_ so that

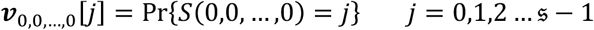

and the matrices 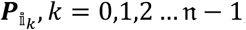; 𝕚 = 𝕕, 𝕙, 𝕡, 𝕣, so that

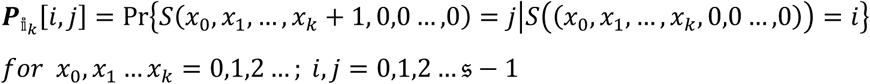

then

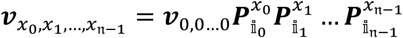

where

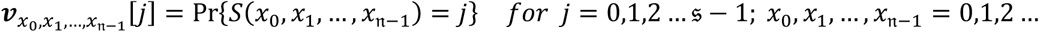

i.e. the 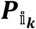 matrices are transition matrices in the same sense that the ***T***_𝕚_ matrices in theorem 6 are, and they are found in the same manner. Hence,

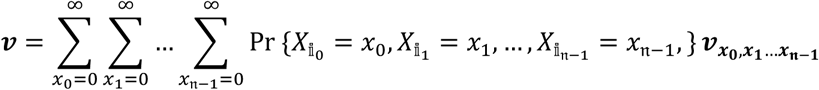

gives the unconditional state distribution. Because unbalanced patterns in anaphase I tetrads are retained in the polar bodies, balanced patterns in anaphase I tetrads are weighted by 2 and unbalanced patterns in anaphase I tetrads are weighted by 0.^3^ So

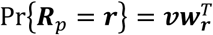

where

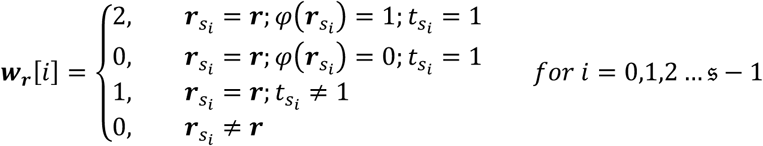

Note that the function

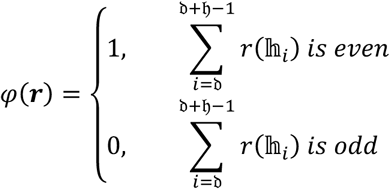

is 1 if the pattern is balanced and 0 if the pattern is unbalanced (see figure 2.5 for an induction proof).

The next step is to find expressions for the probability of observing a given number of chiasma events in each interval. Using an induction argument similar to the one in the proof for theorem 1 gives

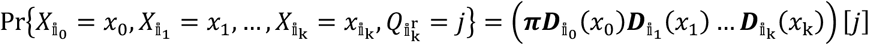

where

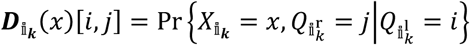

Note that 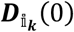 is equivalent to 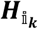 in theorem 1. To get an expression for 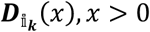, first make the following observations. If there are in total *x* chiasma events in the interval, then between 0 and *x* of them must be *X*′events. If there are *l* (0 ≤ *l* < *x*) *X*′ events in the interval, then we can express the total number of *C* events, *h*, as *h =* (*i O*″events before the first *X*″ event) + (the first *X*″ event) + (*l X*′ events [in no particular spatial order]) + (*n* additional *C*″events up to and including the last *X*″ event) + (the final *q - j O*″ events, where *q* is the phase after the last *X*″ event, so as to end up in phase *j* at the right boundary), in short

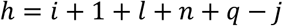

In the special case where *γ*_*m*_ = 1, *n* and *q* are fixed at (*m* + 1)(*x* − *l* − 1) and *m*, respectively, so that we can get an expression for 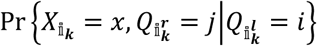 by summing over all possible values of *l*, i.e.

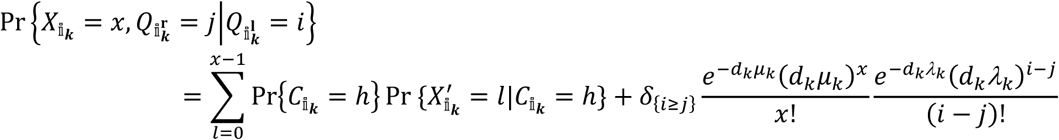

where 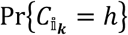 and 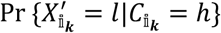 are given by the Poisson and Binomial distributions, respectively (Copenhaver et al 2002). The final term takes into account the case where all the *x* chiasma events are *X*′ events (which means that there are no *X*″ events). This term is non-zero only if *i* ≥ *j* (otherwise there must be at least one *X*″ events, which is a contradiction), and is in that case equal to the probability of observing *x X*′ events and *i* − *j O*″ events, which is just the Poisson probabilities multiplied. For the general model, we have to also sum over all values of *n* and *q*, weighted by their probabilities, i.e.

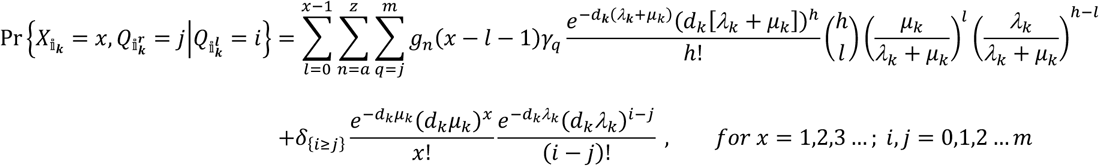

where *a* = *x* − *l* − 1 and *z* = (*x* − *l* − 1)(*m* + 1). The function *g*_*n*_(*s*) gives the probability that the *n-th C*″event after an *X*″ event is the *s-th X*″ event after that *X*″ event. This is achieved by defining the base case *g*_0_(0) = 1, which means that the zeroth *C*″ event is the zeroth *X*″ event with probability 1, and then recursively conditioning on which *X*″ event is the last before the *s-th*, i.e.

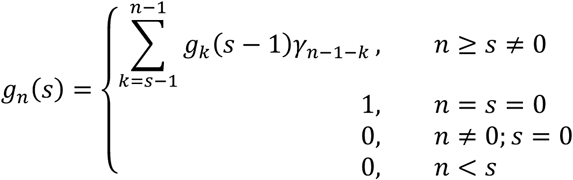

Note the difference between *g*_*n*_(*s*) and *b*_*n*_ from theorem 1: the former gives the probability that the *n-th C*″event after an *X*″ event is strictly the *s-th X*″ event after that *X*″ event, whereas the latter gives the probability that the *n-th C*″event after an *X*″ event is any *X*″ event.

*QED*

## 4 Discussion

I have in this paper suggested a generalization of the many counting models of chiasma interference, and derived novel expressions for recombination pattern probabilities in homo- and heterokaryotypes for two types of chromosomal inversions. I envision two applications for these expressions. Firstly, the model can be useful in testing simplifying assumptions in analytic studies against the results of more realistic numerical simulations. It is, for example, common in multilocus evolution models to simplify the analysis by assuming haploidy and run numerical diploid simulations to check that they give qualitatively similar results (e.g. Kirkpatrick 1982, Kirkpatrick and Servedio 1999). Similarly, such models almost invariably assume no chiasma interference for reasons of mathematical simplicity. Running numerical simulations with various degrees of chiasma interference could prove useful in identifying whether or when this assumption affects the results. This is particularly important in studies of chromosomal inversions, as both recombination rates and underdominance in inversion heterokaryotypes can depend strongly on the degree of interference (Navarro et al. 1997). The model also provides a more realistic view of the relationship between recombination and underdominance in heterokaryotypes. Models of the evolution of inversions commonly assume perfect suppression of chiasmata (e.g. Charlesworth and Charlesworth 1973, Tricket and Butlin 1994, Dagilis and Kirkpatrick 2014), which implies no recombination and no underdominance (equivalent to *d*_*k*_*=0 for all k in the inverted region* in my model). Although some inversions, particulary short ones, seem to have this attribute, it appears that most do not (Coyne et al. 1993, Navarro and Ruiz 1997). Furthermore, some studies specifically designed to test the effect of recombination in heterokaryotypes assume no underdominance (e.g. Feder and Nosil 2009), or vice versa (e.g. Kirkpatrick and Barton 2006), which is unrealistic in that chiasmata that cause recombination will inevitably also cause some degree of sterility, and vice versa. The model presented in this paper thus provides an opportunity to compare the results of simulations with or without the assumption of perfect suppression of chiasmata, and to conduct more realistic simulations of the effects of recombination and underdominance in heterokaryotypes.

Secondly, the predictions of the model with various parameter settings can be compared to real data so as to elucidate problems related to interference and inversions. For example, Lange et al. (1997) perform maximum likelihood analyses using both the pure counting model and a variant of what I call the stochastic counting model to see if the latter gives a significantly better fit to the data. My model allows, for example, for a similar comparison of the two-pathway counting model and the full general model (i.e. of a two-pathway model with deterministic or stochastic counting in the interfering pathway) or of the full general model and the stochastic counting model (i.e. of a stochastic counting model with or without an additional non-interfering pathway). The model can also be used to address questions about interference in inversion heterokaryotypes. I have, for example, throughout this paper assumed that the intermediate events in interval 𝕚_*k*_ are suppressed by a factor *d*_*k*_, and that the counting process works on this depleted set in the same way that it does in homokaryotypes. Whether interference actually works the same way in hetero-as in homokaryotypes is however an open question that can be addressed by comparing the predictions of the model to recombination data. A related question is that of the presence or absence of interference across breakpoint boundaries, which can be addressed by testing the predictions of the model with the parameter *α* set to 1 and 0, respectively.

## 5 Acknowledgements

Figures 1, 4, 5, 6, 7, 8 and 9 were drawn by Sofie Ensby Rostad. I thank Arcadi Navarro, Fabrice Eroukhmanoff, Glenn-Peter Sætre, Thomas Hansen, Åke Brännström and Iain Johnston for discussions, exchanges and comments.

## 7 Appendix: Vector and matrix notation

All vectors and matrices are denoted in bold. A matrix or vector followed by square brackets ([]) denotes the particular element of that matrix or vector, so that e.g.

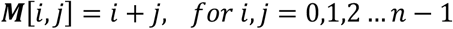

indicate that ***M*** is a zero-indexed matrix of size *n,n* with element *i,j* (row, column) equal to *i*+*j*. All vectors are row-vectors unless otherwise indicated, so that e.g.

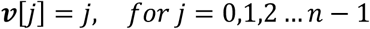

means that

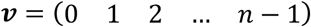

When referring to a column-vector, I either do so by writing out the vector in full, like this:

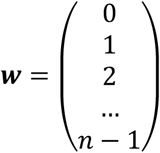

or by first defining the vector elements and then indicate in subsequent expression that the vector is transposed, like this:

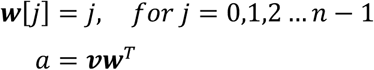

Two vectors or matrices placed side by side always indicate matrix multiplication. When matrix multiplication operations are to be performed on a given number of matrices with increasing indices from left to right, I sometimes write it in the following short form

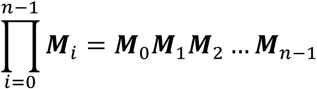

The multiplication of *x* instances of the same matrix (***M***) is denoted ***M***^*x*^, so that

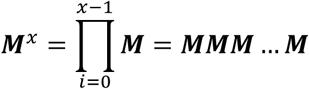

By definition,

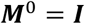

i.e. a matrix to the zeroth power is always equal to the corresponding identity matrix. The *one-vector*, denoted **1**, is a vector of only ones, i.e. **1** = (1 1 1 … 1).

## Footnotes

1 This is sometimes referred to as the *four-strand model of recombination*, to distinguish it from the simplified *two-strand* (sometimes *one-strand*) *model* that is implicit in most textbook accounts (see Speed 1995 for a discussion of these two models). *Strand* is here synonymous with chromatid.

2 The theorem gives the stationary distribution, but any other distribution may be used.

3 If this is not clear, notice that eliminating unbalanced anaphase I patterns is probabilistically equivalent to turning the two unbalanced pattern chromatids of an anaphase I tetrad into copies (one of each) of the two balanced pattern chromatids in the same tetrad, and then drawing randomly (uniformly) from the resulting four balanced pattern chromatids.

